# Functional activation reveals a repressor-gated fourth type VI secretion system in clinical *Pseudomonas aeruginosa*

**DOI:** 10.64898/2026.02.12.705469

**Authors:** Liwen Wu, Yong Liu, Ruolin Huang, Ying Zhao, Junbo Liu, Ying An, Jingtong Su, Yiqiu Zhang, Xingyu Wang, Haoyu Zheng, Xuanling Liu, Tongtong Pei, Xiaoye Liang, Xiaotian Liu, Yu Xia, Min Zheng, Ronghui Liu, Yi Li, Hui Wang, Jiuxin Qu, Yingxia Liu, Liang Yang, Mingjie Zhang, Tao Dong

## Abstract

*Pseudomonas aeruginosa* is a major opportunistic pathogen in which the type VI secretion system (T6SS) contributes to interbacterial competition and virulence. While most strains encode three T6SSs, we here functionally characterize a fourth system (H4-T6SS) in a clinical *P. aeruginosa* isolate. The H4-T6SS gene cluster is transcriptionally silent under laboratory conditions. By promoter rewiring, we demonstrate that the H4-T6SS assembles a contractile machine and mediates effector secretion and robust inter- and intraspecific antagonism. We identify a dominant pore-forming effector and its secretion pathway and a silent yet conserved nuclease effector. Genetic and biochemical analyses show that H4-T6SS expression is activated by an H4-encoded transcriptional regulator and repressed by H-NS family proteins. Analysis of over 1,200 clinical isolates reveals that the H4-T6SS is distributed across lineages and geographic regions. Finally, we observe functional crosstalk between H4- and H2-T6SSs. Together, these findings establish H4-T6SS as a functional and repressor-gated secretion system that expands the competitive arsenal of clinical *P. aeruginosa*.

## Introduction

*Pseudomonas aeruginosa* is a ubiquitous gram-negative bacterium and a leading cause of hospital-acquired infections, particularly in immunocompromised individuals and patients with chronic lung disease ^1–3^. Its remarkable adaptability arises from a large and dynamic accessory genome shaped by horizontal gene transfer and strong selective pressures in polymicrobial environments, generating new traits that enhance survival, persistence, and competitiveness ^4^. Among the most potent weapons encoded by *P. aeruginosa* is the type VI secretion system (T6SS), a contractile nanomachine that delivers toxic effectors directly into neighboring cells ^5^. Structurally analogous to an inverted bacteriophage tail, the T6SS assembles a sheath–tube complex that contracts within milliseconds to propel a spike loaded with effectors across cell envelopes ^6–10^. Known effectors include those targeting cell walls, membranes, nucleic acids, and essential metabolic processes in both prokaryotic and eukaryotic cells ^11–18^.

In *P.* aeruginosa, three functionally independent T6SSs (H1–H3) have been characterized and shown to be tightly regulated to mediate diverse antibacterial, anti-eukaryotic, and community-shaping functions ^19–24^. Although clinical isolates have shown extensive genome variations, most studies of *P. aeruginosa* T6SSs have focused on a small number of laboratory reference strains, most notably PAO1 and PA14. Few newly identified clinical-specific systems have been experimentally validated and the driving mechanisms for such diversity and pathogen evolution remain poorly understood ^25–31^. Recent population-scale genomic analyses reported the existence of a fourth T6SS cluster, designated H4-T6SS, in a subset of *P. aeruginosa* genomes. However, it remains unclear whether it encodes a functional secretion system or a silent genomic relic, how it is regulated, or what effectors it deploys.

Here, we identify and functionally characterize an H4-T6SS in a clinical *P. aeruginosa* isolate obtained from a COVID-19 patient. We show that the H4-T6SS gene cluster encodes all conserved structural components required for secretion but is transcriptionally silenced under laboratory conditions. By promoter engineering, we demonstrate that H4-T6SS assembles dynamically, secretes proteins, and mediates potent antibacterial and anti-eukaryotic activity. We identify H4-T6SS associated effectors and define the function and secretion mechanism of a colicin-like pore-forming effector. We further show the H4-T6SS is regulated by a transcriptional activator and repressed by H-NS family regulators. Extending these findings to a large multi-center collection of clinical isolates, we uncover the prevalence and natural activation potential of H4-T6SS and reveal functional crosstalk with the canonical H2-T6SS. Together, our study establishes H4-T6SS as a potent and tightly regulated secretion system that expands the competitive landscape of clinical *P. aeruginosa* strains.

## Results

### A clinical *P. aeruginosa* isolate encodes a complete H4-T6SS cluster

We isolated a clinical *P. aeruginosa* strain from a COVID-19 patient and identified an additional T6SS gene cluster beyond the canonical H1–H3 systems through genome analysis (NCBI Reference Sequence: NZ_CP061699.1, LYSZa7 strain). This cluster, which we designate H4-T6SS, encodes all conserved structural components required for a functional T6SS, including sheath (TssB/C), tube (Hcp), baseplate, membrane complex, and ATPase (Fig. 1A). Additionally, the H4-T6SS main cluster encodes two VgrG proteins, a PAAR protein, and multiple putative effector genes (Fig. 1A).

**Figure 1.**
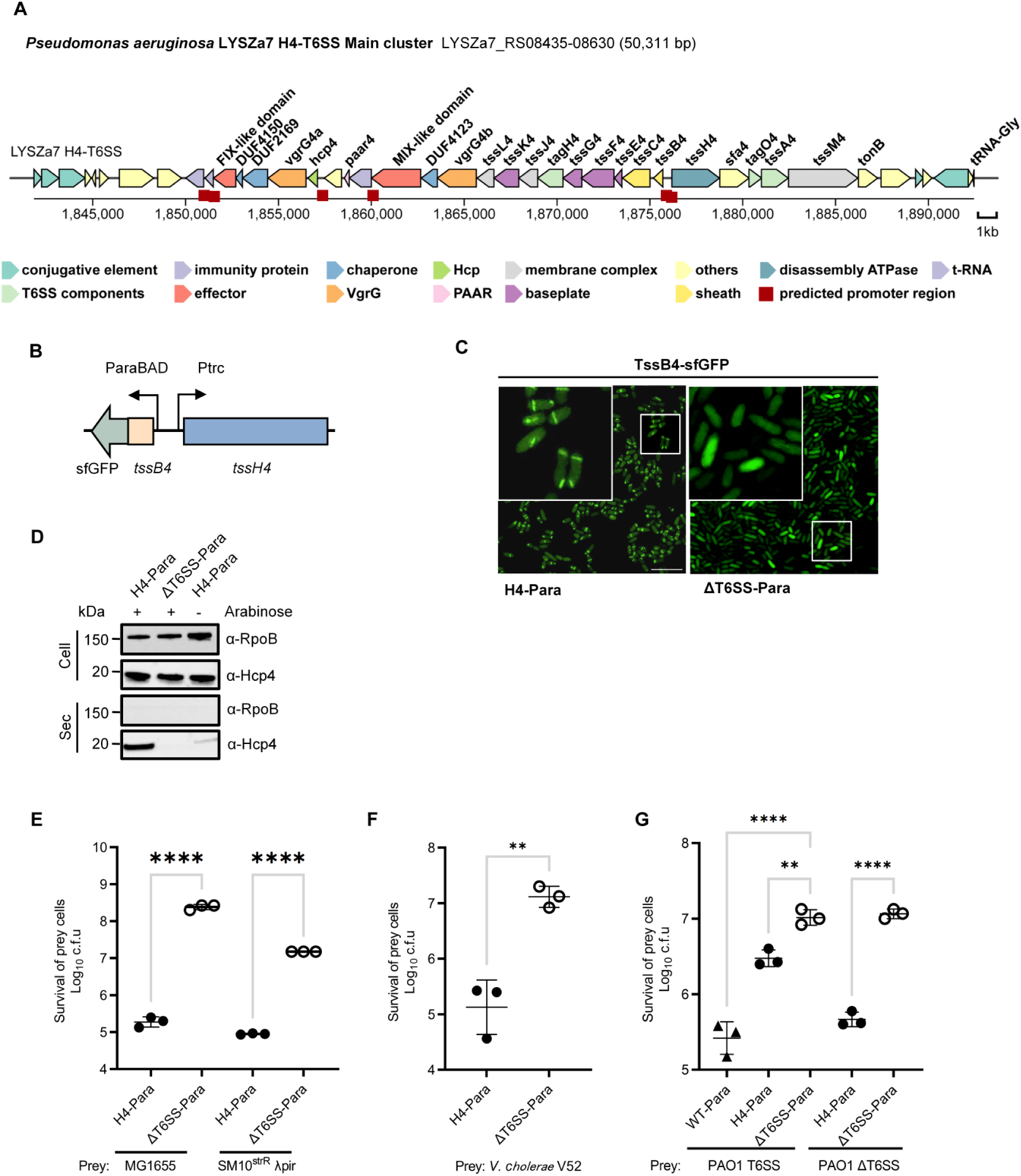
The inducible H4-T6SS exhibits T6SS activity. **A.** Schematic diagram of the H4-T6SS gene cluster in *P. aeruginosa* clinical isolate LYSZa7. Different colored arrows indicate the corresponding protein function category. Red squares indicate the promoter region predicted by Prokaryote Promoter Prediction v2.0. **B.** Genetic modification of the inducible H4-T6SS. The original *tssB4* and *tssH4* promoters were replaced with the arabinose-inducible promoter ParaBAD and the constitutive promoter Ptrc. The TssB4 protein is fused with the sfGFP fluorescent protein. **C.** Fluorescence images showing sheath assembly in H4-Para and ΔT6SS-Para strains. Cells were induced with 0.1% [w/v] arabinose for 3 h during the exponential phase. A representative 30 × 30 μm field of cells with a 3× magnified 5 × 5 μm inset (marked by box) is shown. Scale bar: 5 μm. **D.** Secretion analysis of Hcp4 in H4-Para and ΔT6SS-Para strains. Cells were grown at 37°C for 4 h to OD_600_ ∼1, with or without 0.1% [w/v] arabinose. Competition assays of the H4-Para and ΔT6SS-Para against **E.** *E. coli* MG1655 and SM10^strR^ λpir, **F.** *V. cholerae* V52 and **G.** *P. aeruginosa* PAO1 strains. For **E**-**G**, cells of killer and prey were mixed at a ratio of 10:1 or 20:1 (killer: prey), and co-incubated for 12 h at 30 °C. Error bars indicate the standard deviation of three biological replicates. Statistical significance was calculated using a Student’s t-test (**E** and **F**) and a one-way ANOVA test (**G**) for each group, **P < 0.01, ****P < 0.0001.

To assess whether the H4-T6SS is functional in LYSZa7, we constructed a strain in which the essential structural genes of the H1–H3 systems were inactivated (Δ*tssM1*Δ*tssM2*Δ*tssB3*), thereby isolating H4-T6SS activity (designated H4). A further deletion of *tssM4* generated a T6SS-null strain (ΔT6SS), which served as a negative control. For detection of H4-T6SS secretion, we raised a polyclonal antibody against the H4-specific Hcp protein (Hcp4) (Fig. S1A). Under laboratory conditions (aerobic growth in LB medium), Hcp4 was not expressed intracellularly or secreted into the culture supernatant (Fig. S1B). Consistent with this observation, time-lapse fluorescence microscopy and killing assays failed to detect assembly of the H4 sheath protein TssB4 under conditions known to activate other *P. aeruginosa* T6SSs, including low-temperature growth or RP4-mediated conjugation from a *Escherichia coli* donor ^32,33^ (Fig. S1C,D). These results indicate that although the H4-T6SS gene cluster encodes a complete secretion apparatus, it remains inactive under the tested conditions.

To further examine this inactivity, we fused three predicted promoters within the H4-T6SS main cluster to a luciferase reporter (Fig. 1A, Fig. S1E). When introduced into *E. coli*, the promoters of *tssB4* and *tssH4* drove detectable reporter expression, whereas no expression was observed in *P. aeruginosa* PAO1 or LYSZa7. The *hcp4* promoter was inactive in both *E. coli* and *P. aeruginosa* (Fig. S1E). These data suggest that H4-T6SS promoters are subject to repression in *P. aeruginosa*. Together, these results indicate that the H4-T6SS is transcriptionally silenced in *P. aeruginosa*.

### Induced activation reveals that H4-T6SS is functional

To test whether H4-T6SS is capable of assembly and secretion when expressed, we rewired its regulatory architecture by replacing the native promoter upstream of *tssB4* with an arabinose-inducible promoter (ParaBAD) and the promoter upstream of *tssH4* with a constitutive Ptrc promoter ^34^ (Fig. 1B). This strain (H4-Para) is expected to drive coordinated expression of the left and right arms of the H4-T6SS gene cluster. A T6SS-null strain carrying the same promoter replacements (ΔT6SS-Para) served as a control.

Upon arabinose induction, fluorescence microscopy revealed robust assembly of H4-T6SS sheaths, and Western blot analysis confirmed intracellular expression and secretion of Hcp4 in the H4-Para strain but not in the ΔT6SS-Para control (Fig. 1C,D; Video S1). Note that the Hcp4 antibody is specific because no signal was detected in the Δ*hcp4* deletion mutant (Fig. S2A). Reversing the orientation of the inducible and constitutive promoters also supported H4-T6SS assembly (Fig. S2B). In contrast, inducing either arm of the cluster alone was insufficient to trigger assembly, consistent with the requirement for coordinated expression of both operons (Fig. S2B). We next examined whether the activated H4-T6SS confers competitive advantages. In interspecies competition assays, induction of H4-T6SS resulted in efficient killing of *E. coli* strains MG1655 and SM10, as well as *Vibrio cholerae* V52 (Fig. 1E,F; Fig. S2C,D). Note that SM10 and V52 but not the MG1655 strain are known to induce the post-translational activation of H1-T6SS in *P. aeruginosa* reference strain PAO1 ^33,35^. In intraspecies competition, the H4-Para strain outcompeted PAO1, with enhanced killing observed against a PAO1 T6SS-null mutant (Fig. 1G; Fig. S2E). Moreover, induction of H4-T6SS caused cytotoxicity toward the murine macrophage cell line RAW264.7 (Fig. S2F). These results demonstrate that H4-T6SS is a fully functional secretion system capable of targeting both bacterial and eukaryotic cells when transcriptionally activated.

### Secretome analysis identifies H4-T6SS–dependent effector proteins

To define the effector repertoire of the H4-T6SS, we performed comparative secretome analysis of the induced H4-Para strain and the ΔT6SS-Para control. Liquid chromatography–tandem mass spectrometry (LC–MS/MS) analysis of culture supernatants identified 1,682 proteins in the H4-Para strain and 1,451 proteins in the ΔT6SS-Para strain. Seventy-five proteins were significantly enriched in the H4-Para secretome (Fig. S3A). Among these, we detected Hcp4, VgrG4a, VgrG4b, and peptides corresponding to a putative effector encoded by RS08520 (Fig. 2A). These results indicate that induction of H4-T6SS leads to secretion of both core structural components and associated effector proteins.

**Figure 2.**
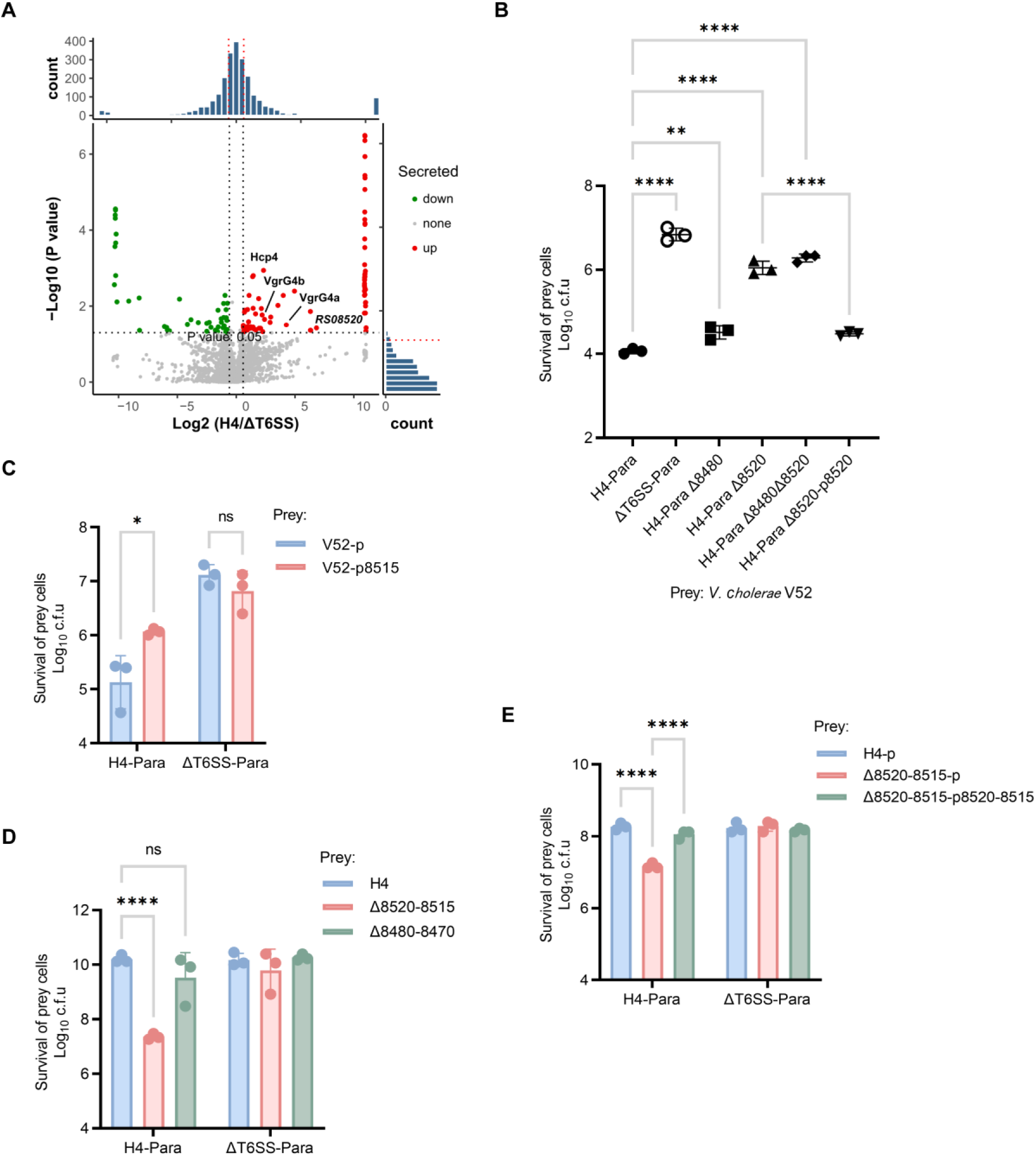
RS08520 is a dominant antibacterial effector of H4-T6SS. **A.** Volcano plot showing the log 2 ratio of secreted proteins in the H4-Para and ΔT6SS-Para strains to the corresponding p-value (−Log 10 ). Each red dot represents a significantly high abundance protein (up) after the comparison of H4-Para and ΔT6SS-Para. Each green dot represents a significantly low abundance protein (down). And each gray dot represents a non-significantly altered protein (none). The samples for mass spectrometry analysis were the induced supernatant samples shown in Figure 1D. **B.** Interspecific competition between H4-Para effector deletion mutants and *V. cholerae* V52. **C.** Competition assay of the H4-Para and ΔT6SS-Para against *V. cholerae* V52 complemented with an empty vector (pPSV37) or a vector expressing the immunity protein 8515. **D.** Intraspecific competition between the H4-Para and ΔT6SS-Para strains and effector-immunity deletion mutants Δ8480-8475&8470 and Δ8520-8515. **E.** Competition assay of the H4-Para and ΔT6SS-Para against the effector-immunity deletion mutant Δ8520-8515 complemented with an empty vector (pPSV37) or a vector expressing the immunity protein 8515. For **B-E**, the proteins carried by the pPSV37 plasmid were induced by 1 mM IPTG during both the liquid culture and co-incubation processes. Cells of killer and prey were mixed at a ratio of 10:1 or 20:1 (killer: prey), and co-incubated for 12 h at 30 °C. Error bars indicate the standard deviation of three biological replicates and statistical significance was calculated using a one-way ANOVA test, *p < 0.05, **p < 0.01, ****p < 0.0001, ns, not significant.

### RS08520 is essential for H4-T6SS–mediated antibacterial activity

To determine the contribution of individual effectors to H4-T6SS function, we generated deletion mutants lacking RS08520. In interspecies competition assays against *V. cholerae*, deletion of RS08520 abolished H4-T6SS–dependent killing (Fig. 2B; Fig. S3B). Expression of the downstream gene RS08515, predicted to encode an immunity protein, protected *V. cholerae* from H4-mediated killing (Fig. 2C; Fig. S3C).

In intraspecies competition assays, a mutant lacking RS08515 was susceptible to killing by the H4-Para strain, and this phenotype was rescued by complementation with RS08515 (Fig. 2D,E; Fig. S3D,E). These findings strongly suggest that RS08520 is a bona fide H4-T6SS effector that confers antibacterial activity and that RS08515 functions as its cognate immunity protein.

### RS08520 is a pore-forming toxin delivered by a cognate chaperone–VgrG–PAAR pathway

Sequence analysis revealed that RS08520 belongs to the BTH_I2691 family and contains an N-terminal MIX (Marker for Type VI effectors) domain ^36^, a central region homologous to nuclear pore proteins, and a C-terminal colicin-like pore-forming domain (Fig. 3A). To assess its toxicity, RS08520 was expressed in the periplasm of *E. coli* via a twin-arginine translocation signal ^37^. Periplasmic expression resulted in severe growth inhibition, which was neutralized by co-expression of RS08515 (Fig. 3B; Fig. S3F). Mutational analysis showed that disruption of predicted transmembrane helices within the C-terminal domain significantly reduced toxicity (Fig. 3B), suggesting that membrane insertion is critical for effector function. Purified GFP-tagged RS08520 bound to liposomes *in vitro* and formed ion-conducting pores in planar lipid bilayers, whereas a C-terminal truncation mutant lacked pore-forming activity (Fig. S4A-C). Together, these data establish RS08520 as a pore-forming effector.

**Figure 3.**
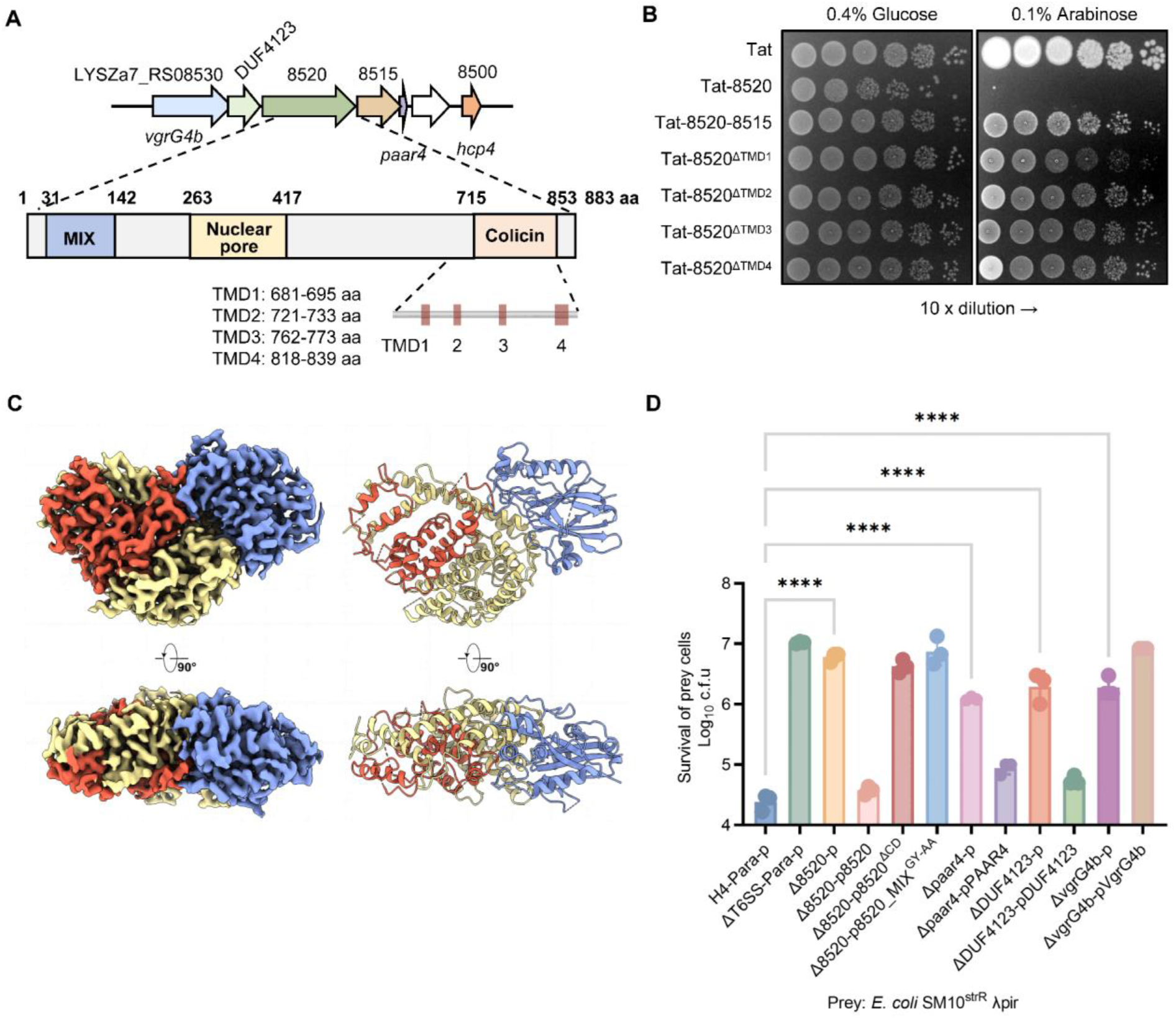
Structural and functional characterization of RS08520. **A.** Prediction of the RS08520 conserved domain and function. The conserved functional domains and transmembrane regions were predicted using HHPred and DeepTMHMM. **B.** Bacterial toxicity assay of expressing RS08520 with its downstream immunity protein and expressing RS08520 TMD mutants in *E. coli* periplasm. *E. coli* cells carrying the pBAD24 vector were plated on 0.1% [w/v] arabinose and 0.4% [w/v] glucose plates for protein induction and repression, respectively. **C.** Front and bottom views of the cryo-EM density map of RS08520 (Left). Atomic model of RS08520 in ribbon representation (Right). NTD, the central linker and CTD are colored in blue, yellow and red, respectively. **D.** Competition assay showing the effects of RS08520 and its upstream/downstream mutants on H4-Para interspecific competition. The GY-AA mutant carries mutations G88A and Y91A in the conserved GxxY motif of the MIX domain^36^. The ΔCD mutant lacks the CTD 715-853 amino acids. The proteins carried by the pPSV37 plasmid were induced by 1 mM IPTG during both the liquid culture and co-incubation processes. Cells of killer and prey were mixed at a ratio of 10:1 (killer: prey), and co-incubated for 12 h at 30 °C. Error bars indicate the standard deviation of three biological replicates and statistical significance was calculated using a one-way ANOVA test, ****p < 0.0001.

### Cryo-EM structure and AlphaFold 3 analyses reveal two conformations of RS08520

Despite the presence of predicted membrane-interacting elements, RS08520 behaved as a soluble protein and was purified to homogeneity in the absence of detergents (Fig. S5A). Single-particle cryo-electron microscopy yielded a structure at 3.0 Å resolution (Fig. 3C; Supplemental Table 2), allowing atomic modeling of most of the polypeptide chain, with the exception of a short N-terminal segment and a central flexible linker. The structure reveals a compact three-domain architecture. Notably, hydrophobic elements required for membrane insertion are sequestered within the protein core.

In addition, a fraction of the protein partitioned into membrane fractions during purification, suggesting structural plasticity (Fig. S5B). To explore this further, we used AlphaFold 3 to probe alternative conformational states. Independent prediction runs converged on two major conformations: a compact soluble state closely resembling the cryo-EM structure, and an extended state in which hydrophobic C-terminal residues are exposed (Fig. S5C-E). These observations support a model in which RS08520 exists in equilibrium between a soluble state and a membrane-inserted pore-forming state, consistent with its periplasmic toxicity and pore-forming activity.

Despite extensive efforts, we were unable to isolate a stable RS08520–RS08515 complex experimentally. AlphaFold 3 modeling predicts that RS08515 binds directly to the C-terminal toxin domain of RS08520, shielding hydrophobic residues implicated in membrane insertion (Fig. S5F). Such binding is stabilized by complementary electrostatic interactions, including negatively charged residues at the base of the RS8520 cleft (Asp277, 842; Glu297, 843) and corresponding positively charged residues on the immunity protein (Arg283, 289; Lys287) (Fig. S5F).

### RS08520 is delivered by a dedicated chaperone–VgrG–PAAR pathway

Genetic analysis showed that deletion of the upstream DUF4123 chaperone, the cognate VgrG4b, or PAAR4 abolished H4-T6SS–dependent killing, without affecting T6SS assembly or secretion (Fig. 3D; Fig. S6A,B; Video S2). Mutations in the MIX domain or deletion of the C-terminal toxin domain similarly eliminated effector activity. Complementation restored killing in all cases except *vgrG4b*, likely reflecting stoichiometric constraints.

To understand how RS08520 is recruited to the secretion apparatus, we used AlphaFold 3 to model the quaternary effector–chaperone–spike complex. The predicted structure reveals that RS08520 forms a stable subcomplex with its chaperone that engages the C-terminal extension of the VgrG spike, while PAAR caps the spike tip (Fig. S6C). In this configuration, the PAAR-sharpened VgrG punctures the target envelope, enabling translocation of RS08520 via peripheral cargo loading. Consistent with this model, deletion of *paar* significantly impaired RS08520-mediated bactericidal activity (Fig. 3D; Fig. S6A).

Altogether, these findings suggest that RS08520 is a dominant H4-T6SS toxin delivered via a VgrG–chaperone–PAAR-dependent pathway.

### RS08480 encodes a conserved H4-associated nuclease

Unlike RS08520 that is only found in the LYSZa7 strain, another putative effector RS08480 is highly conserved among H4-positive clinical isolates (Fig. S7A). Bioinformatic analysis predicted RS08480 to contain a FIX (Found in type sIX effector) domain and encode a nuclease-like protein ^38^ (Fig. 4A). Consistent with this prediction, expression of RS08480 in *E. coli* resulted in DNA degradation and growth inhibition, which was neutralized by co-expression of a downstream immunity gene (Fig. 4B–D). However, RS08480 was not robustly detected in H4-dependent secretome analyses under inducible conditions, and deletion of RS08480 had minimal impact on H4-T6SS–mediated antibacterial competition (Fig. 2A,B,D; Fig. S3B,D). These results suggest that RS08480 is a conserved and yet conditionally active nuclease effector whose contribution to H4-T6SS function may depend on specific regulatory or environmental contexts.

**Figure 4.**
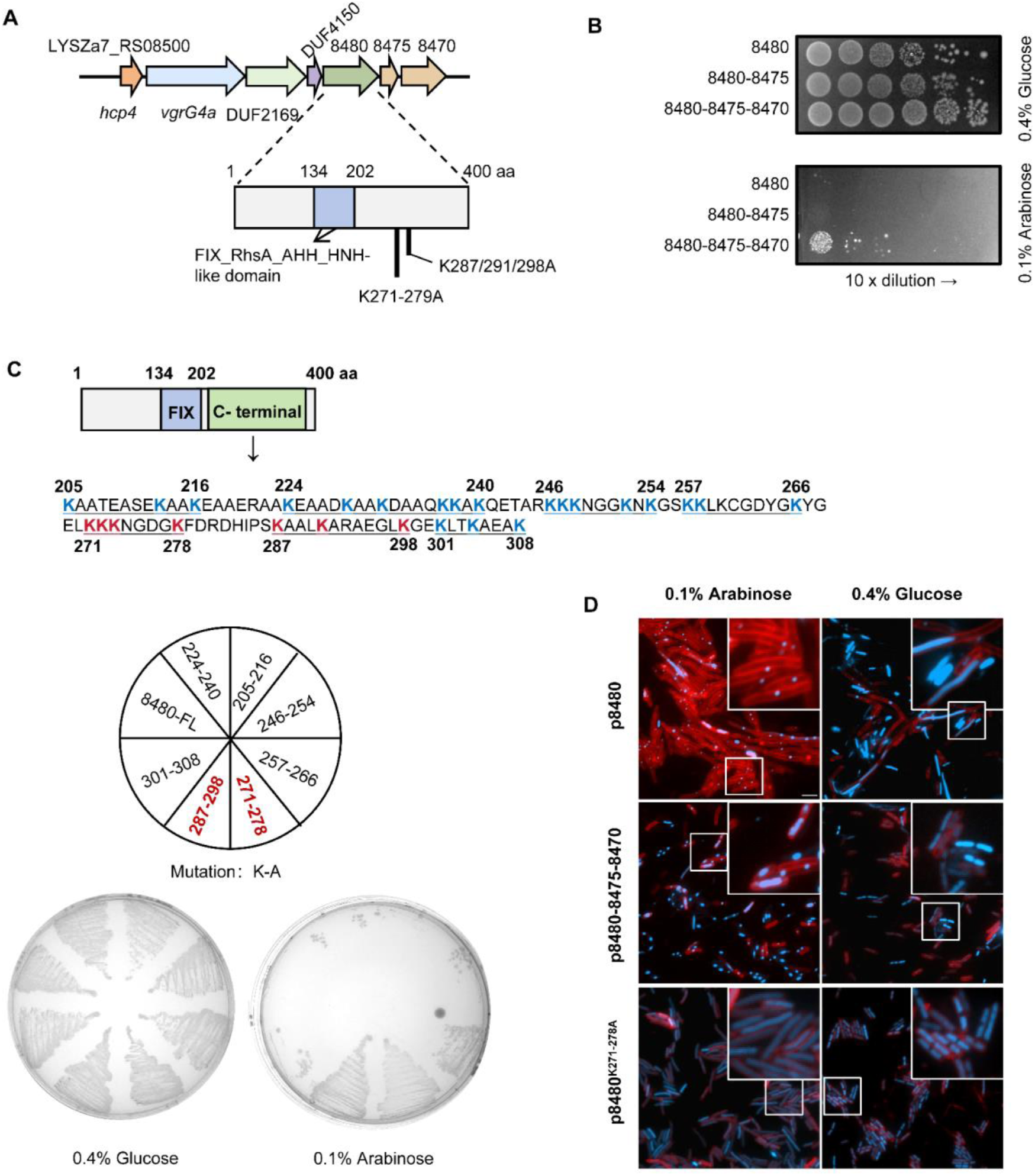
RS08480 is a conserved effector with nuclease activity. **A.** Prediction of the RS08480 conserved domain. The conserved functional domain was predicted using HHPred and CD-Search. **B.** Bacterial toxicity assay of the effector RS08480 and its downstream immunity proteins when targeting *E. coli* cytoplasm. *E. coli* cells carrying the pBAD24 vector were plated on 0.1% [w/v] arabinose and 0.4% [w/v] glucose plates for protein induction and repression, respectively. **C.** Toxicity assay of the RS08480 C-terminal mutants. The RS08480 mutation sites indicated in the sector correspond to the mutations of lysine combinations in each underscore of the C-terminal amino acid sequence above. All lysines were mutated to alanines. *E. coli* cells carrying the pBAD24 vector with full-length RS08480 protein (8480-FL) and mutant proteins were cultured on 0.1% [w/v] arabinose and 0.4% [w/v] glucose plates for protein induction and repression, respectively. **D.** Fluorescence microscopy images showing the nucleic acid and cell length alterations of *E. coli* cells expressing RS08480 full length protein, immunity proteins and the attenuated mutant. Cells were grown to the exponential phase and induced with 0.1% [w/v] arabinose or repressed with 0.4% [w/v] glucose for 1 h at 37°C. Nucleic acid (blue) and inner membrane (red) were stained by 0.5 μg/mL DAPI and 5 μg/mL FM4-64 for 20 min. A representative 30 × 30 μm field of cells with a 3× magnified 5 × 5 μm inset (marked by box) is shown. Scale bar: 5 μm.

#### H4-T6SS encodes a Fis-family transcriptional activator

The H4-T6SS gene cluster encodes a predicted Fis-family transcriptional regulator Sfa4 (WP_200200388.1) located downstream of *tssH4*, sharing homology with the RpoN-dependent activators Sfa2 and Sfa3 that regulate the H2- and H3-T6SSs ^39^. To test whether this regulator activates H4-T6SS expression, we employed transcriptional reporters for the *tssB4*, *tssH4* and *hcp4* promoter (Fig. S1E; Fig. S8A). Ectopic expression of the H4-associated Sfa4 protein in *E. coli* robustly activated the *hcp4* reporter only, suggesting that the *hcp4* promoter is controlled by Sfa4 (Fig. S1E). Additionally, deletion of *sfa4* abolished Hcp4 expression in the inducible H4-T6SS strain and the associated competition (Fig. S8B,C). Although ectopic expression of Sfa4 also activated Hcp4 expression in the wild type *P. aeruginosa* strain, no H4-T6SS-dependent competition was observed (Fig. S8B,C).

These results suggested that Sfa4 is a positive regulator for Hcp4 expression and that H4-T6SS expression is constrained by dominant repression in *P. aeruginosa*.

### H-NS–like proteins MvaT and MvaU repress H4-T6SS

To identify factors mediating H4-T6SS repression, we performed DNA pull-down assays using H4 promoter regions followed by mass spectrometry. This approach consistently identified the H-NS-like protein MvaT as a candidate repressor (Fig. 5A; Fig. S9A). Deletion of *mvaT*resulted in partial and heterogeneous activation of H4-T6SS reporters, whereas deletion of *mvaU*, encoding another H-NS-like paralog, had little effect (Fig. S9B). Electrophoretic mobility shift assays further demonstrated that MvaT bind directly to H4-T6SS promoter regions (Fig. S9C), supporting a model in which H4-T6SS is subject to xenogeneic silencing by H-NS-like repressors. Notably, simultaneous deletion of *mvaU* and partial inactivation of *mvaT* (using an *mvaT* truncation allele, as the double deletion was not viable in our hands) led to significant activation of *tssB4* and *tssH4* transcription, as measured by quantitative PCR, and enabled detectable H4-T6SS expression signals by fluorescence microscopy (Fig. 5B-E). Complementation with a single or two proteins can largely suppress the majority of the activation signal (Fig. 5C-E). We further show that H4-T6SS promoters are relatively AT-rich compared with the genomic average (Fig. S9D), a known feature of loci targeted by MvaT/MvaU-mediated repression ^40^.

**Figure 5.**
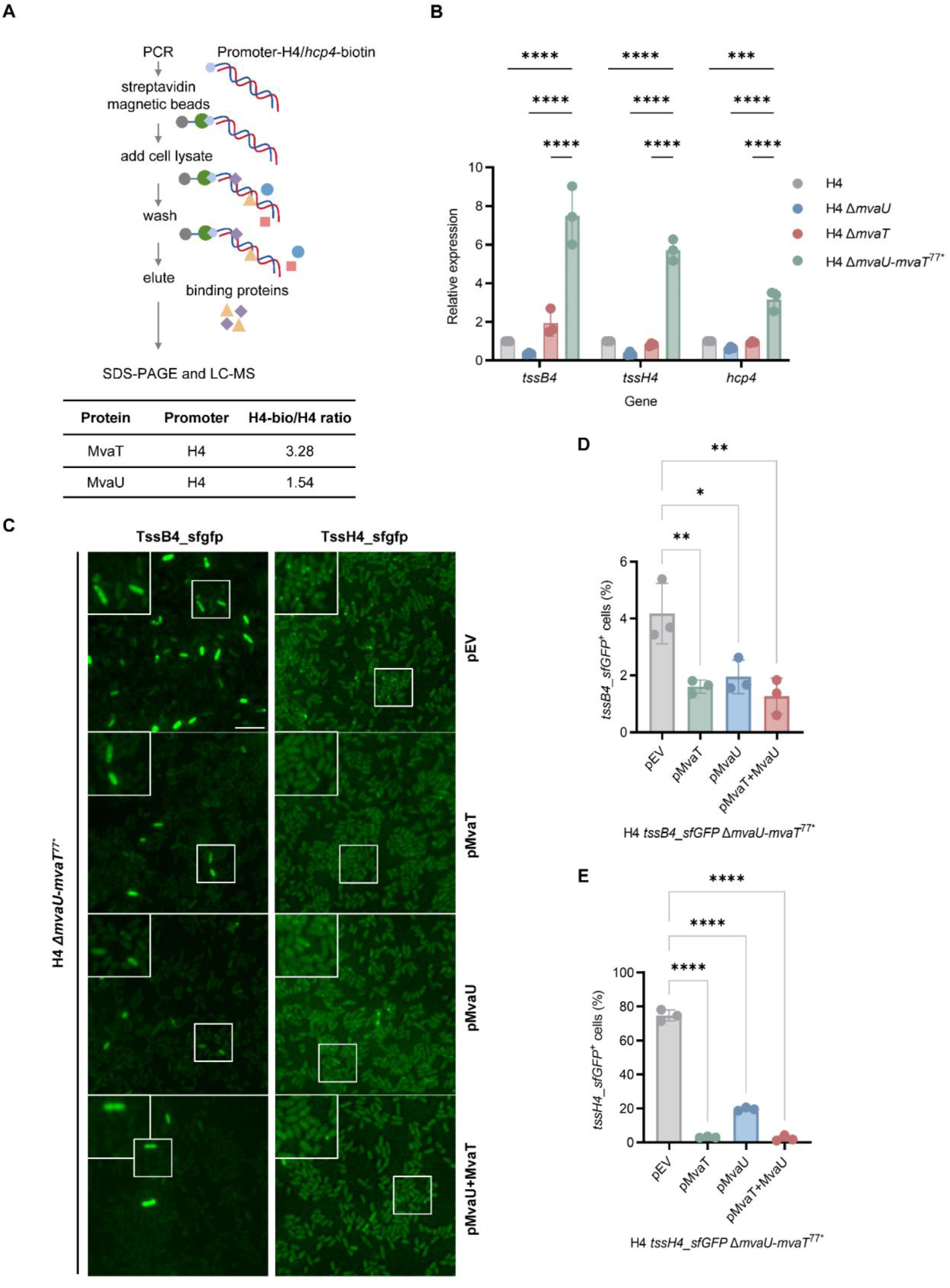
H-NS family regulators repress H4-T6SS promoters. **A.** Method flowchart of DNA pull-down assay. Cell lysates were incubated with each DNA probe, captured with streptavidin beads and analyzed by SDS-PAGE and LC-MS. The un-biotinylated H4 and *hcp4* promoters were used as the negative control. By calculating the ratio of the protein abundance pulled down by the biotinylated DNA probe to the non-specifically bound protein abundance, hit proteins were identified. **B.** Quantitative real-time PCR analysis of H4 gene expression. Samples run in triplicate with SYBR green detection. Ct values were normalized to 16SrRNA gene and analyzed by the ΔΔCt method. Error bars indicate the standard deviation of three biological replicates and statistical significance was calculated using a one-way ANOVA test, ****p < 0.0001. **C.** Fluorescence images showing sfGFP signals of TssB4 and TssH4 in H4 Δ *mvaU*-*mvaT*^77^* mutant and MvaT/MvaU-complementation strains. A single protein was induced using 1 mM IPTG. Two proteins were induced simultaneously using 1 mM IPTG and 0.1% [w/v] arabinose. A representative 30 × 30 μm field of cells with a 3× magnified 5 × 5 μm inset (marked by box) is shown. Scale bar: 5 μm. Proportion of cells forming chromosomally encoded TssB4_sfGFP (**D**) and TssH4_sfGFP_(**E**) signals in H4 Δ*mvaU*-*mvaT*^77^* mutant and MvaT/MvaU-complementation strains. For **D** and **E**, error bars represent the mean ± standard deviation calculated from three distinct 30 × 30 μm fields of view, with a total of at least 800 cells analyzed. Statistical significance was calculated using one-way ANOVA test for each group; ****p < 0.0001.

Together, these results indicate that H4-T6SS expression is gated by xenogeneic silencing and requires relief of MvaT/MvaU repression in addition to the presence of a cognate Fis-family activator.

### H4-T6SS is present in diverse clinical isolates and shows evidence of natural activation

To assess the prevalence of H4-T6SS, we screened approximately 1,200 *P. aeruginosa* clinical isolates collected in Shenzhen and 94 isolates from Beijing using H4-specific PCR primers (Fig. 6A; Fig. S10A). Twenty-nine isolates (2.2%) were positive for H4-T6SS (Fig. 6B). Dot blot analysis identified several clinical isolates with detectable Hcp4 expression, suggesting native activation (Fig. 6C). The complete H4-T6SS gene cluster of one H4-T6SS positive strain C2 was determined by whole genome sequencing. Apart from differences in the effector types, the T6SS core structure components were near identical to those of the LYSZa7 strain (Fig. 6D). These findings indicate that the H4-T6SS is distributed across multiple clinical lineages and may be naturally active under specific conditions.

**Figure 6.**
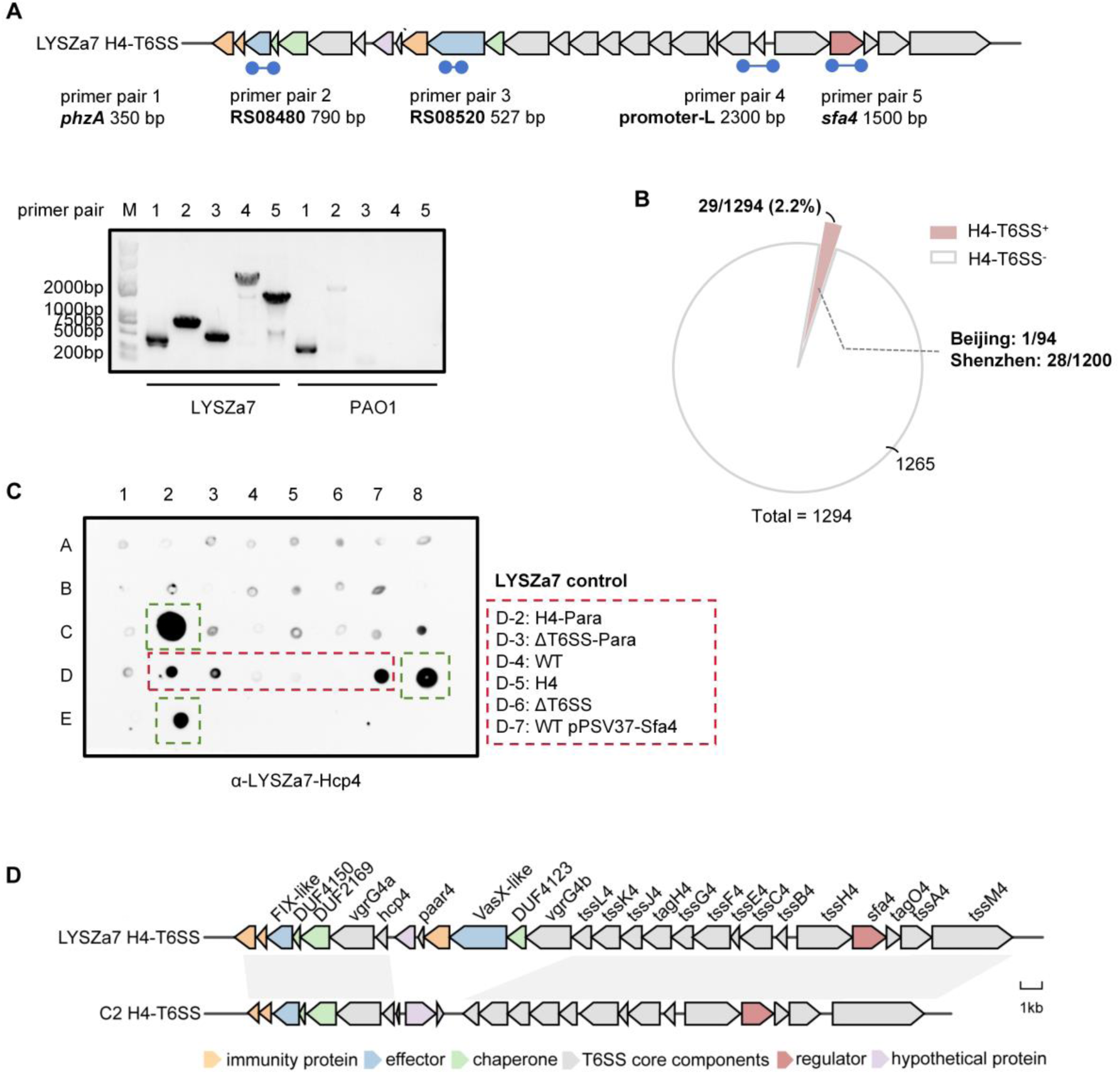
Screening of H4-T6SS in *P. aeruginosa* clinical isolates. **A.** Specificity detection of the H4-T6SS primers in LYSZa7 and PAO1. *phzA* is used as a positive gene for identifying the *Pseudomonas* genus. The blue circles indicate the regions amplified by different primer pairs. M: marker; primer pair 1-5: the primer pair used for each lane’s PCR product. **B**. Prevalence of H4-T6SS genetic elements in *P. aeruginosa* clinical strains (n=1294). The pink area represents the proportion of H4-T6SS positive strains. **C.** Dot blot for detecting the Hcp4 expression in H4-T6SS positive strains from Figure 6B (total 28 clinical isolates). The red dashed box indicates the position of the LYSZa7 control samples. The green dashed box indicates the possible activated *P. aeruginosa* clinical isolates. Samples are positioned by both horizontal numbers and vertical letters. **D.** Schematic diagram of the H4-T6SS gene cluster in *P. aeruginosa* clinical isolate C2 from Figure 6C. The gray shading between the two gene clusters represents the gene regions with similar functions.

### Crosstalk between H4-T6SS and H2-T6SS

Upon ectopic expression of TssB4 in the LYSZa7 wild-type strain that is H4-T6SS inactive, we unexpectedly observed a 100-fold increase of T6SS-mediated antibacterial activity against *E. coli*, suggesting functional crosstalk between distinct T6SSs (Fig. 7A; Fig. S11A). Live-cell imaging of LYSZa7 and of PAO1, which lacks an H4-T6SS cluster, revealed the formation of sheath-like structures upon expression of TssB4–sfGFP, indicating that TssB4 can be recruited into a heterologous T6SS assembly (Fig. 7B; Video S3). Systematic analysis using a panel of T6SS mutants in both LYSZa7 and PAO1 demonstrated that these sheath-like structures formed only in the presence of an intact H2-T6SS, indicating H2-T6SS as the recipient for TssB4 incorporation (Fig. 7B; Fig. S11B). Reciprocally, in the H4-Para strain in which only the H4-T6SS is present and activated, ectopic expression of TssB2–sfGFP supported sheath assembly, indicating bidirectional compatibility between TssB2 and TssB4 (Fig. 7C; Fig. S11C).

**Figure 7.**
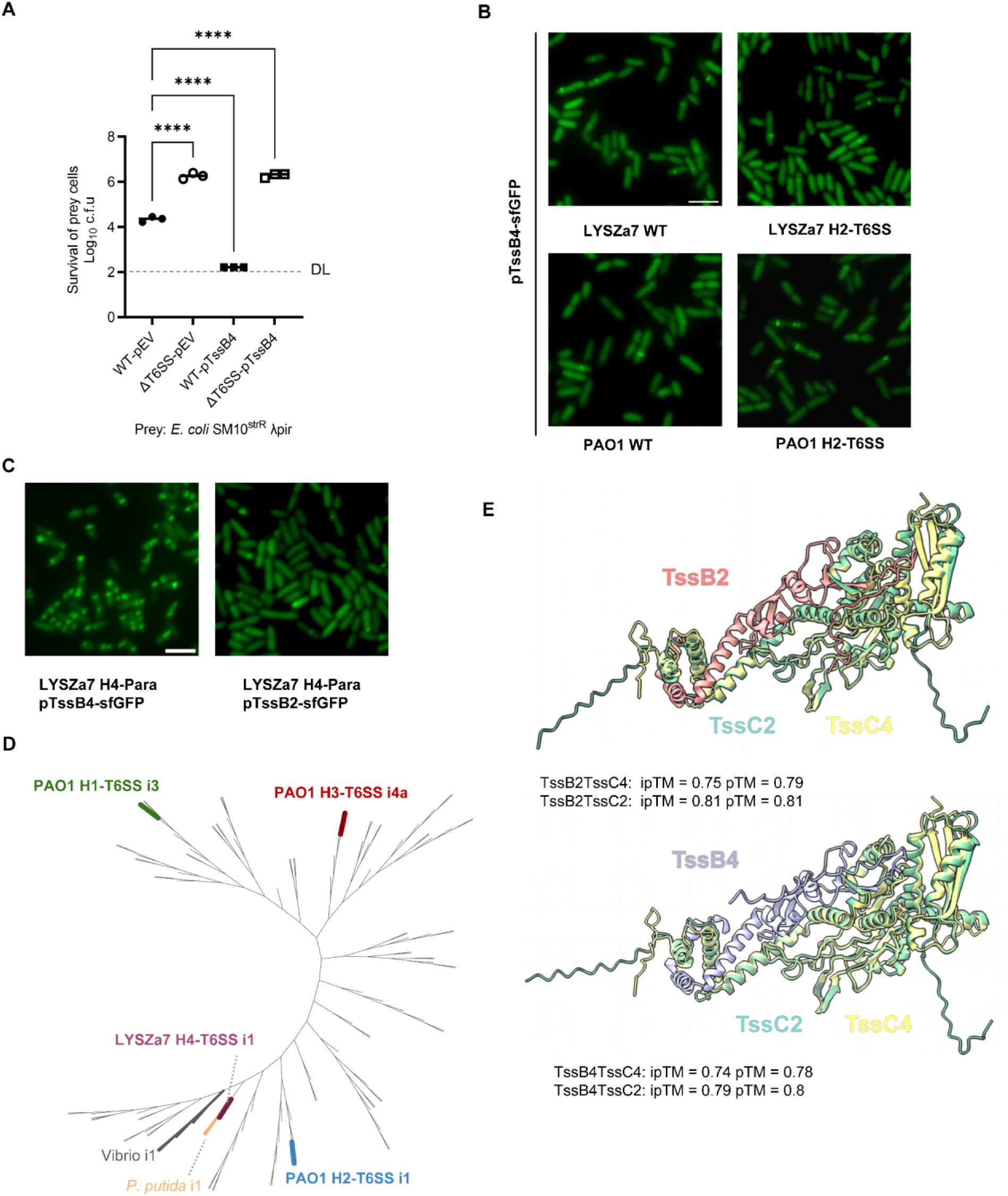
Crosstalk occurs between H4-T6SS and H2-T6SS. **A.** Competition analysis between LYSZa7 and *E. coli* prey. Wild type and H1-H4-inactivated LYSZa7 cells carrying the empty plasmid vector pPSV37 or TssB4 were used as killer cells. An *E. coli* SM10 derivative strain was used as the prey. Competition was performed on solid LB plate for 24 h at 28°C and both killer and prey cells were enumerated by serial dilutions on selective media. DL, detection limit. Error bars indicate the standard deviation of three biological replicates and statistical significance was calculated using a one-way ANOVA test, ****p < 0.0001. **B, C**. Fluorescence microscopy images showing the assistance of TssB4 in the assembly of H2-T6SS and the assistance of TssB2 in the assembly of H4-T6SS. TssB4 and TssB2 proteins carrying by pPSV37 vector were induced to express using 1mM IPTG. A representative 15 × 15 μm field of cells is shown. Scale bar: 3 μm. **D.** Classification of TssB4. TssB4 protein sequence of LYSZa7 strain was classified using SecReT6. Different colors represent the branches where PAO1 TssB1-3 and LYSZa7 TssB4 are located and the types of T6SSs. The genera that exhibit similar evolutionary relationships to LYSZa7 are also marked in color. **E.** Alphafold 3-predicted the interaction between TssB2/TssB4 and TssC2/TssC4. TssB2, TssB4, TssC2 and TssC4 are colored in red, purple, green and yellow, respectively.

**Figure 8.**
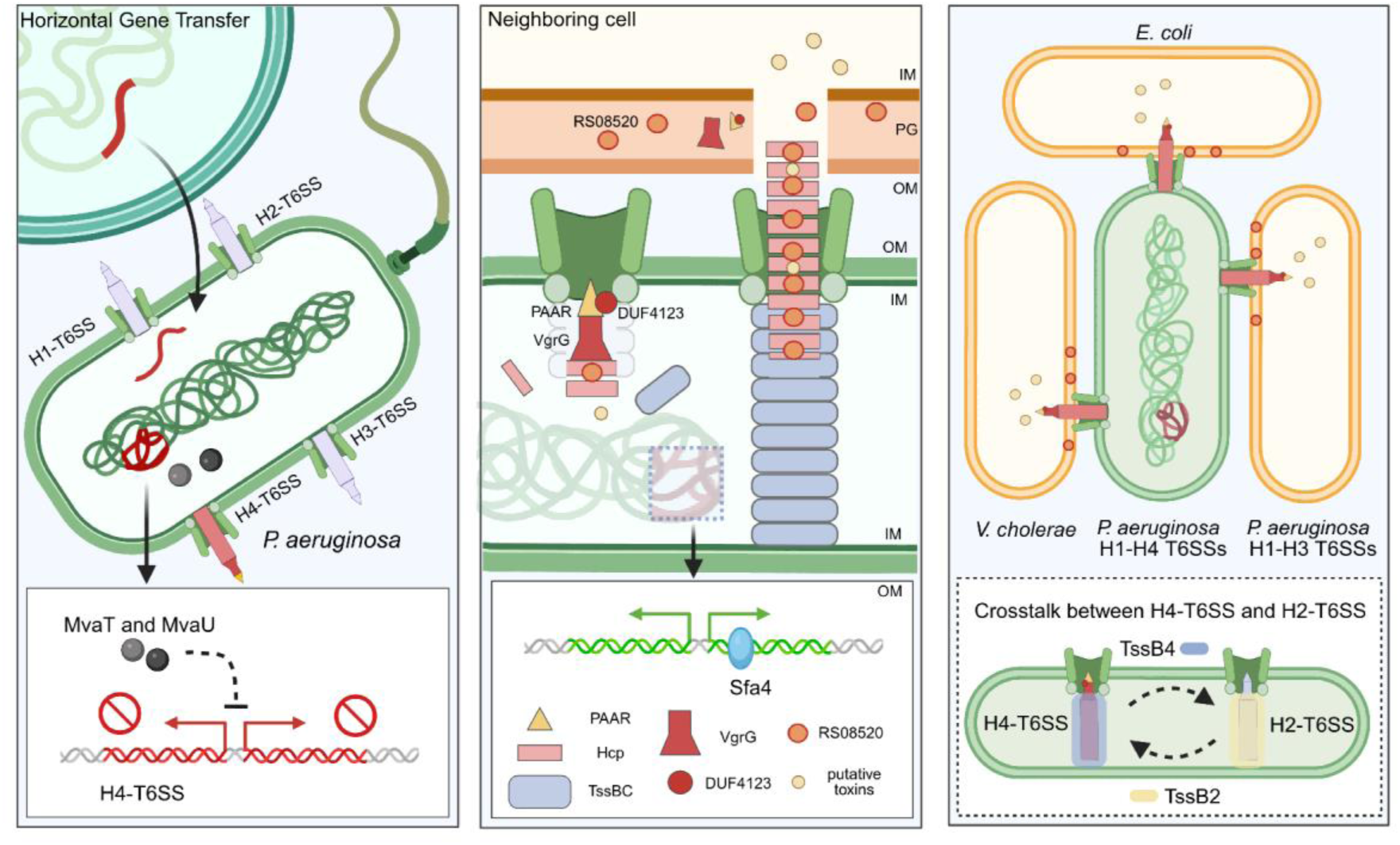
Schematic representation illustrating the regulation and assembly of the H4-T6SS. Left panel: The H4-T6SS gene cluster, potentially acquired via horizontal gene transfer, remains silenced in *P. aeruginosa* LYSZa7 strain by H-NS-like repressors MvaT and MvaU. Middle panel: Upon activation, the H4-T6SS assembles into a functional apparatus capable of effector delivery into neighboring cells. Sfa4 is a H4-T6SS-encoded positive regulator for Hcp expression. Right panel: Activated H4-T6SS confers inter- and intra-species competitiveness to the *P. aeruginosa* clinical isolate. During assembly, functional crosstalk occurs between H2-T6SS and H4-T6SS.

To further test whether the observed sheath exchangeability reflects an underlying structural compatibility between the H2- and H4-T6SS, we examined their evolutionary relatedness and the predicted interactions between their core sheath components. Phylogenetic analysis classified the H4-T6SS as an i1-type T6SS, closely related to the H2-T6SS (Fig. 7D) ^41–43^. Consistent with this evolutionary proximity, AlphaFold 3 modeling predicted that heterologous TssB–TssC pairs (TssB2–TssC4 and TssB4–TssC2) can form stable hybrid complexes with high confidence, providing a structural basis for the interoperability observed between H2- and H4-T6SS components (Fig. 7E).

Together, these results demonstrate that TssB2 and TssB4 are functionally interchangeable and can be incorporated into heterologous T6SS assemblies, providing direct evidence for structural crosstalk between co-existing T6SSs.

## Discussion

T6SS was first defined in 2006 through parallel work in *V. cholerae* and *P. aeruginosa*, establishing these organisms as founding models for T6SS research. Subsequent landmark discoveries in *P. aeruginosa* revealed that T6SSs function as antibacterial weapons through toxin–immunity pairs and can engage in dynamic “dueling” and retaliatory “tit-for-tat” behaviors during interbacterial encounters ^35^. Here, we extend this lineage of discoveries by establishing that a fourth T6SS cluster, previously identified only through comparative genomics, encodes a functional and regulated secretion apparatus in clinical *P. aeruginosa*. Although H4-T6SS encodes all conserved structural components, it is repressed under the laboratory conditions tested, underscoring that genomic completeness alone does not imply activity. By promoter rewiring, we show that H4-T6SS assembles as a canonical contractile machine, secretes effectors, and confers strong competitive advantages, establishing it as a conditionally deployed nanoweapon in a subset of clinical isolates.

Repression of the H4-T6SS is consistent with the broader principle that *P. aeruginosa* T6SSs are differentially regulated and frequently silenced. Although the H4-T6SS locus encodes a Fis-family activator for *hcp4* transcription, this activity is masked by repression mediated by the H-NS–like proteins MvaT and MvaU. Such behavior is characteristic of horizontally acquired, AT-rich loci in *P. aeruginosa* and other gram-negative bacteria where xenogeneic silencing constrains potentially costly or disruptive functions until appropriate conditions are met. Known conditions that relieve H-NS mediated silencing include changes in DNA supercoiling, osmolarity, temperature, surface association, quorum sensing, and the action of anti-silencing factors ^44–51^. The H4-T6SS may be activated by one of those general signals or a specific signal to H4-T6SS. Alternatively, it is also possible that the H4-T6SS represents a relatively recent acquisition that has not yet been fully integrated into host regulatory networks, rendering genetic screening unfeasible. Future research is required to distinguish these scenarios by identifying native conditions that alleviate repression and determining whether H4-T6SS activation is transient, heterogeneous, or coordinated with other stress and competition responses. Nonetheless, this regulatory logic places H4-T6SS within the broader framework of conditionally deployed accessory systems, in which silencing and activation are layered to balance evolvability with cellular homeostasis.

An unexpected and conceptually important finding of this study is the functional interoperability between the H4- and H2-T6SSs, considering the high specificity in protein-protein interactions required for assembly. Although *P. aeruginosa* encodes multiple T6SS clusters that are typically considered discrete and independently regulated, we find that sheath components can be shared between H4- and H2-T6SSs, enabling cross-system assembly. To our knowledge, such crosstalk between distinct T6SS clusters have not been demonstrated. Additionally, detecting such interoperability between coexisting T6SSs is challenging. Here, the use of fluorescently labelled sheath proteins provides a valuable and convenient readout of cross-system interactions. These observations also raise the possibility that additional forms of crosstalk involving non-labelled structural elements or regulatory factors may have been overlooked. Interoperability could introduce layers of coordination, competition, or interference between secretion systems, to either potentiate or constrain activity depending on cellular context. Understanding such interactions, especially between existing systems and a newly acquired T6SS, will be an important future direction for deciphering how multi-T6SS bacteria deploy these systems in complex and fluctuating environments.

When derepressed, H4-T6SS in LYSZa7 delivers a potent pore-forming toxin RS08520. Notably, this effector is strain-specific and poorly conserved, highlighting that not only the T6SS machineries but also their cargo effectors are evolving among clinical strains. We have also identified an auxiliary effector island flanked by mobile genetic elements elsewhere in the genome that is not present in other H1-H3 systems (Fig. S7B). Therefore, although the H4-T6SS is currently rare among clinical isolates, its functionality, conditional deployment, and association with mobile genetic elements indicate that *P. aeruginosa* continues to diversify its competitive strategies in clinical settings. Whether such systems directly influence disease severity or treatment outcomes remains difficult to estimate at present. Nevertheless, the emergence and retention of additional secretion systems highlight an evolutionary trajectory toward increasingly complex interbacterial interactions and the need for continued surveillance of *P. aeruginosa* in clinical environments.

## Author contributions

T.D. conceived the project. L.W., Y.L., R.H., Y.Z., J.L., Y.A., J.S., Y.Z., X.W., H.Z., X.L., T.P. and X.L. performed research. M.Z., L.Y., Y.L., J.Q., H.W., Y.L., R.L., M.Z., Y.X., and X.L. provided key reagents and materials. T.D. and L.W. wrote the manuscript.

## Acknowledgments

This work was supported by funding from Shenzhen Medical Research Fund (B2402028), National Natural Science Foundation of China (W2431022, 3253000254, W2433062, 32400096), the Guangdong Innovative and Entrepreneurial Research Team Program (2023ZT10Y013), Guangdong basic and applied basic research foundation (2024A1515010319), Science and Technology Program of Shenzhen (KCXFZ20230731100901003) and Shenzhen Key Laboratory of Biochip (grant number SYSPG20241211173949065). The funders had no role in study design, data collection, interpretation, or the decision to submit the work for publication.

## Materials and Methods

### Bacterial strains and plasmids

Strains and plasmids are listed in Supplementary Table 1, respectively. Strains were grown in LB (1% [w/v] tryptone, 0.5% [w/v] yeast extract, 0.5% [w/v] NaCl) aerobically at 37 ℃ unless otherwise stated. Antibiotics were used as follows: gentamycin (50 µg/ml), carbenicillin (100 µg/ml), streptomycin (300 µg/ml), irgasan (25 µg/ml). Precise gene deletion and insertion mutants were generated via homologous recombination using the pEXG2.0 suicide vector. LYSZa7 proteins expressed in *E. coli* were cloned onto the pBAD24 or pET28a vectors, while the proteins expressed in *P. aeruginosa* (Pa) and *V. cholerae* were cloned onto the pPSV37 vector. All constructs were confirmed by sequencing.

### Bacterial competition assay

Killer cells were grown in liquid LB to OD_600_ ∼ 1 and prey cells were collected from overnight LB-agar plates. Cells were harvested by centrifugation at 12,000 × *g* for 2 min and resuspended in fresh LB. For Pa intraspecies competition, killer cells and prey cells were mixed at a ratio of 20:1 (killer:prey) and spotted on LB-agar plates for 12 h at 30°C. For the interspecies competition with *E. coli* and *V. cholera*, killer cells and prey cells were mixed at a ratio of 10:1 (killer:prey) and spotted on LB-agar plates for 12 h at 30°C. After co-incubation, cells were resuspended in 500 µL fresh LB. Survival of killer and prey cells was quantified by serial dilution and plating on LB-agar plates with selective antibiotics. For the competition assays of inducible H4-T6SS strains, 0.1% [w/v] arabinose was added during both the liquid culture and co-incubation processes for induction. For the competition assays of protein complementation, 1 mM IPTG was added during both the liquid culture and co-incubation processes for induction. Error bars show standard deviation of three biological replicates.

### Western blot analysis and dot blot analysis

For western blot analysis, proteins were separated by an SDS-PAGE gel (Yeasen Biotechnology) and transferred to a PVDF membrane (BioRad). For dot blot analysis, Pa cells were collected from overnight LB-agar plates and normalized to OD_600_ = 5 in ddH_2_O. Cells were heated to 98°C for 5 min, and 5 μL of each sample was spotted onto a PVDF membrane as dots and air-dried for 15-30 min. Membranes were blocked with 5% [w/v] non-fat milk in TBST (50 mM Tris, 150 mM NaCl, 0.1% [v/v] Tween 20, pH 7.6) for 1 h at room temperature, then incubated overnight at 4 °C with primary antibodies in antibody diluent buffer (1% [w/v] non-fat milk, 50 mM Tris, 150 mM NaCl, 0.1% [v/v] Tween 20, pH 7.6). After washing, the membranes were incubated with HRP-conjugated secondary antibodies for 1 h at room temperature. Bands were detected by the Clarity Western ECL substrate (Bio-Rad). Monoclonal antibody was ordered from Biolegend (Product # 663905 [RpoB]). The polyclonal antibody to Hcp4 (LYSZa7_RS08500) was customized by Shanghai Youlong Biotech. Secondary antibodies were ordered from ZSGB-Bio (Product #ZB-2305 [mouse] and # ZB-2301 [rabbit]).

### Protein secretion assay

Cells were cultured in 10-20 mL liquid LB at 37°C to OD_600_ ∼1. After centrifugation at 8,000 × *g* for 15 minutes, the supernatants were collected by filtering through 0.22 μm filter. Pellets were resuspended in SDS-loading buffer (Epizyme) and used as whole-cell samples. Supernatants were precipitated in TCA (trichloroacetic acid, 10% [v/v]) at 4°C overnight and centrifuged at 16,000 × *g* for 50 min at 4°C. Pellets were then resuspended in Tris buffer (1.5 M Tris, pH 8.8). Whole-cell and secretion samples were denatured at 98°C for 10 min prior to SDS-PAGE and Western blot analysis. For the secretion of inducible H4-T6SS strains, 0.1% [w/v] arabinose was added into the liquid culture. For secretome analysis, 50 mL supernatants of arabinose-induced cultures were precipitated and loaded to a SDS-PAGE gel. Gel slices containing secreted proteins were cut out and sent for LC-MS/ MS analysis performed by ShenZhen SMQ Group Medical Laboratory.

### Protein toxicity assay

*E. coli* cells carrying different plasmids were grown on LB-agar plates with 0.4% [w/v] glucose at 30°C overnight, and then collected and normalized to OD_600_ = 1 in fresh liquid LB. A ten-fold serial dilution was plated on LB-agar plates containing 0.1% [w/v] arabinose or 0.4% [w/v] glucose for protein induction and repression, respectively. Each experiment was conducted in at least two independent repeats, with one representative experiment shown.

### Fluorescence microscopy

For LYSZa7 and PAO1 strains, cells were grown in liquid ^LB^ to OD_600_ ∼1 at 37°C. The cells were then concentrated to OD_600_∼10 and spotted onto 1% agarose-0.5 × PBS pads. Images were acquired using the Olympus IX85 P1ZF microscope with a 100× oil objective lens (NA 1.5) and a laser of 480 nm. Fiji software was used to process images.

### DNase assay *in vivo*

*E. coli* cells carrying the pBAD24 plasmids for expression of RS08480 full length protein, variants and immune proteins were grown in liquid LB and induced or repressed with 0.1% [w/v] arabinose or 0.4% [w/v] glucose. The cultures were then normalized to OD_600_∼10 in 1 × PBS and stained with 0.5 mg/mL 49, 6-diamidino-2-phenylindole (DAPI; Thermo Scientific) and 5 mg/mL FM4-64 (Thermo Scientific) for 15 min at room temperature. Samples were spotted onto 1% agarose-0.5 × PBS pads. Images were acquired using a Zeiss Axio Observer7 inverted microscope with a 60× oil objective lens (NA 1.4). DAPI (353 nm) and FM4-64 (473 nm) were used for fluorescence excitation. ZEISS ZEN 3.8 software was used to process the images.

### Protein purification

His-tagged proteins were expressed using the pET28a vectors in *E. coli* BL21(DE3). The cells were grown in liquid LB to OD_600_∼0.6 at 37°C. The cultures were then induced with 1 mM IPTG at 16°C for 18 h. The cells were subsequently harvested by centrifugation at 4,500 × *g* for 15 min. The pellets were resuspended in lysis buffer (20 mM Tris-HCl, 150 mM NaCl, 10 mM imidazole, pH 8.0) with 2 mM phenylmethylsulfonyl fluoride (PMSF; Beyotime) and lysed by French press. Lysates were centrifuged at 15,000 *× g* for 40 min and the supernatants were incubated with Ni-NTA resin (Smart-lifesciences). Proteins were eluted in elution buffer (20 mM Tris-HCl, 150 mM NaCl, and variable concentrations of imidazole, pH 8.0). Eluted samples were analyzed by SDS-PAGE analysis. For the liposome binding assay, the *n*-dodecyl-β-D-maltopyranoside (DDM) was added for the protein extraction and purification. Cell lysates were centrifuged at 3,000 *× g* for 10 min, and then 1.5% [w/v] DDM was added to the supernatants for protein extraction at 4°C for 1 h. Insoluble debris was removed by centrifugation 150,000 *× g* for 1 h, and the supernatant was incubated with Ni-NTA beads. For the Ni-NTA affinity, proteins were eluted in elution buffer (20 mM Tris-HCl, 150 mM NaCl, 0.02% [w/v] DDM and variable concentrations of imidazole, pH 8.0).

### Cytotoxicity assay

Host cells were inoculated in 24-well plates at a density of 5×10^5^ per well in DMEM containing 10% FBS overnight. The next day, cells were washed with 1×PBS, and then PBS was replaced with fresh antibiotic-free DMEM containing 10% FBS. LYSZa7 cells were cultured in liquid LB with 0.1% [w/v] arabinose at 37°C to OD_600_∼1. Bacteria cells (MOI 50) were added into wells containing host cells. Cells were washed after 2 h incubation, then incubated with DMEM containing 100 μg/mL gentamycin and 10% PS for 2 h to kill remaining extracellular bacteria, after which they were washed again. After the final wash, 0.2 μg/mL propidium iodide (PI; Sigma-Aldrich) was added to the media for IncuCyte experiments.

### Liposome preparation

*E. coli* total lipids dissolved in chloroform were dried under a stream of nitrogen and the resulting lipid film was vacuum desiccated overnight in order to remove any remaining organic solvent. Lipid films were re-suspended in Tris buffer (20 mM Tris, 150 mM NaCl, pH 7.5) to a final concentration of 25 mg/mL. For the protein binding assay, the lipid solution was diluted in incubation buffer (20 mM Tris, 150 mM NaCl, pH 7.5) to a final concentration of 2.5 mg/mL with extensive pipetting and vortexing for about 10 min. Then, liposomes were prepared via sonication (60 W, 50 % duty cycle, 5 min in total). The solution was centrifuged at 5,000 × *g* for 15 min to remove large aggregates.

### Liposome binding assay

For liposome binding, 5 μM GFP and GFP-RS08520 protein purified with or without DDM was mixed with 2 mg/mL liposome in incubation buffer (20 mM Tris, 150 mM NaCl, pH 7.5, −/+ 0.02% DDM) at a 40 μL volume. After half an hour of incubation at room temperature, the 40 μL mixture was ultracentrifuged at 550,000 × *g* for 30 min at 4℃ to precipitate all liposomes. The supernatant was collected as sample S, and the pellet was resuspended in 40 μL of the 1 x SDS-loading buffer as sample P. Supernatant and pellet samples were analyzed by SDS-PAGE with Coomassie blue staining.

### DNA pull-down assay

PCR was used to amplify biotinylated and non-biotinylated H4 and *hcp*4 promoters as DNA probes. The amplified DNA fragments were purified and incubated with 50 μL streptavidin magnetic beads (Sangon) at room temperature for 30 min. The beads were then washed once each with NEBuffer™ 4 (50 mM Potassium Acetate, 20 mM Tris-acetate, 10 mM Magnesium Acetate, 1 mM DTT, pH 7.9) and B/W buffer (10 mM Tris–HCl pH 7.5, 1 mM EDTA, 2 M NaCl). LYSZa7 cells were grown in liquid LB to OD_600_∼0.8 at 37°C. The cells were then collected by centrifugation, resuspended in ^l^_ysis buffe_^r^ (20 mM Na_2_HPO_4_, 5 mM β-mercaptoethanol, pH 7.0), and lysed by sonication on ice. After centrifugation, the acquired supernatant containing protease inhibitors was mixed with the DNA fragments immobilized on the beads and incubated at room temperature for 1 h. Subsequently, the beads were washed three times with the B/W buffer to remove non-specific proteins. The bound proteins were resolved by SDS-PAGE and stained with the silver staining kit (Sangon). Differential protein bands excised from the SDS-PAGE gel were sent for LC-MS/ MS analysis performed by ShenZhen SMQ Group Medical Laboratory.

### RNA extraction and qRT-PCR analysis

Cells were cultured on LB-agar plate. The cells were then lysed in lysis buffer (1% [w/v] SDS, 2 mM EDTA) and hot acidic phenol (65℃) at 65℃ in a heat block for 5 min, after which the tubes were placed on ice for 10 min. The top supernatant obtained by centrifugation was mixed with an equal-volume of absolute ethanol and precipitated overnight at −20℃. RNA was subsequently harvested by centrifugation at 13,000 × *g* for 15 min at 4℃ and washed twice with 75% ethanol. Genomic DNA was removed by DNase I (NEB) treatment at 37℃ for 30 min. Purified RNA was electrophoresed on 1% [w/v] agarose gel to monitor the integrity and contaminants. Reverse transcription was performed using HiScript III All-in-one RT SuperMix Perfect for qPCR (Vazyme). qRT-PCR reaction was prepared by ChamQ Blue Universal SYBR qPCR Master Mix (Vazyme) and detected by Q9600 Series Real-Time PCE System (BIO-GENER). The 16S rRNA gene was used as the reference. Analysis of relative gene expression was calculated using the 2^−ΔΔCT^ method. Each sample was measured in triplicate and repeated at least three times.

### Electrophoretic mobility shift assays (EMSA)

The P*H4*, P*hcp4* and P*algD* DNA fragments were amplified by PCR using specific primers (Supplementary Table 4). DNA probe (50 ng) was incubated with increasing concentrations of purified His-tagged MvaT protein in a 20 μL reaction (10 mM Tris, 50 mM KCl, 1 mM dithiothreitol, and 0.4% glycerol, pH 7.5). After incubation for 30 min at 28 °C, the samples were electrophoresed on a 6% nondenaturing acrylamide gel in 0.5 × TBE (Tris-borate-EDTA) buffer (44.5 mM Tris base, 44.5 mM boric acid, 1 mM EDTA, pH 8.5) on ice at 120 V for 2 h. The gel was then stained in 0.5 × TBE buffer containing 10,000-fold-diluted CelRed nucleic acid dye (Goyoobio), and DNA was visualized under UV illumination.

### Luciferase assay

The Lux reporter plasmids were constructed using the *tssB4*, *tssH4* and *hcp4* promoters fused with the luciferase enzyme. *E. coli* and *P. aeruginosa* cells carrying the luciferase reporter plasmids were plated on LB-agar plate. After incubation at 37°C for 6 h, the bioluminescent signal was detected by chemiluminescence.

### Pore-forming activity in lipid bilayers

The two compartments of a Teflon chamber were physically separated by a 30 μm-thick Teflon film with a ca. 100 μm orifice where the bilayer formed. The orifice was pre-treated with 1% [v/v] hexadecane in pentane and air-dried thoroughly. Electrolyte buffer (20 mM Tris-HCl, 150 mM NaCl, pH 8.0) was added to both compartments and covered the orifice. A pair of freshly prepared Ag/AgCl electrodes were placed on both sides of the chamber, in contact with the electrolyte buffer. The side that is electrically grounded is defined as the *cis* side, whereas the opposite side is defined as the *trans* side. 1,2-Diphytanoyl-sn-glycero-3-phosphocholine (DPhPC; Avanti Polar Lipids) was added to the electrolyte buffer. After lipid bilayer formation, protein was added to the *cis* compartment. All single-channel recordings were performed by an Axopatch 200B patch clamp amplifier and digitally sampled by a USB-6003 (National Instruments) at a sampling frequency of 50 kHz and low-pass filtered with a corner frequency of 10 kHz. The recorded current traces were filtered with a 2.5 kHz low-pass Bessel filter. Data were analyzed using MATLAB.

### Cryo-EM sample preparation, data collection, and processing

Cryo-EM sample preparation and data collection were performed using modified versions of previously described procedures [PMID:40512840]. A 3.5 μL aliquot of purified RS08520 protein (1 mg/mL) was applied to a freshly glow-discharged Quantifoil holey carbon grid (R 1.2/1.3, 300 mesh; Quantifoil, Jena, Germany). Grids were blotted for 4.5 s at ∼100% humidity and plunge-frozen in liquid ethane using a Vitrobot (Thermo Fisher Scientific).

Cryo-EM data were collected at liquid nitrogen temperature on a Titan Krios transmission electron microscope (Thermo Fisher Scientific) operated at 300 kV and equipped with a K3 direct electron detector (Gatan). Movies were recorded in super-resolution counting mode with a calibrated physical pixel size of 0.92 Å (0.46 Å in super-resolution mode). The dose rate was set to 25 electrons per physical pixel per second. Each movie had a total exposure time of 1.46 s, resulting in an accumulated dose of 50 electrons/Å² fractionated into 32 frames. All movies were recorded with a defocus range of 1.0 to 2.0 μm. Detailed data collection parameters are listed in Supplementary Table 2.

For data processing, motion correction and dose weighting were performed using the EMshark implementation of MotionCor2 ^52^, then imported into cryoSPARC for contrast transfer function (CTF) estimation using Patch CTF Estimation^53^. Particles were picked using a combination of Blob Picker, Template Picker, and Topaz, followed by extraction for two-dimensional classification. False positives and poor-quality particles were removed through several rounds of 2D classification. An initial three-dimensional model was generated using Ab-Initio Reconstruction. Further cleaning was performed using Heterogeneous Refinement, and the best classes were subjected to Non-Uniform Refinement with C1 symmetry to enhance resolution. The overall resolution was determined using the gold-standard Fourier Shell Correlation (FSC) = 0.143 criterion. Local resolution estimates were generated using cryoSPARC.

The initial atomic model was generated using AlphaFold 3 and docked into the cryo-EM map using UCSF Chimera ^54,55^. Manual model building and adjustment were performed in Coot ^56^. The model was improved through iterative rounds of manual adjustment in Coot and real-space refinement using PHENIX ^57^. All structure figures were prepared using UCSF ChimeraX ^58^.

### Bioinformatic analysis

All gene sequences of *P. aeruginosa* strain LYSZa7 were retrieved from the draft genome assembly (GenBank NZ_CP061699.1), managed and analyzed by Benchling. The RS08520 and RS08480 sequence was analyzed with the CD-search tool of NCBI webserver ^59^, HHPred interactive server ^60^ and DeepTMHMM webserver. The classification of T6SS was based on the maximum likelihood phylogenetic tree of TssB homologs. The phylogenetic tree contains the TssB4 and 126 experimentally validated TssB proteins recorded in SecReT6 ^61^. Phylogenetic analysis is performed using MAFFT v7.475, FastTree v2.1.11 and visualized by Interactive Tree Of Life (iTOL). The predicated monomer and complex structures of RS08520 was generated by AlphaFold 3. The homologous gene sequences of RS08480 were retrieved by BLASTN, and then aligned using NCBI Multiple Sequence Alignment Viewer, Version 1.26.0.

