## Supplementary Table 1 and Supplemental Table 2 for "Functional activation reveals a repressor-gated fourth type VI secretion system in clinical *Pseudomonas aeruginosa*"

**Supplementary Table S1. Strains and plasmids used in this study.**

| Bacterial strains | Genotype | Description | Source |
|---|---|---|---|
| ***Pseudomonas aeruginosa* LYSZa7** | WT | clinical isolate from Department of Clinical Laboratory, Shenzhen Third People's Hospital, The Second Affiliated Hospital of Southern University of Science and Technology, Shenzhen, Guangdong, China | |
| | *ΔtssM24tssB3* | H1; in-frame deletion of *LYSZa7_RS14210, LYSZa7_RS08600 and LYSZa7_RS17780* in LYSZa7 WT | This study |
| | *ΔtssM14tssB3* | H2; in-frame deletion of *LYSZa7_RS00480, LYSZa7_RS08600 and LYSZa7_RS17780* in LYSZa7 WT | This study |
| | *ΔtssM124* | H3; in-frame deletion of *LYSZa7_RS00480, LYSZa7_RS14210, LYSZa7_RS08600 and LYSZa7_RS17780* in LYSZa7 WT | This study |
| | *ΔtssM12tssB3* | H4; in-frame deletion of *LYSZa7_RS00480, LYSZa7_RS14210,and LYSZa7_RS17780* in LYSZa7 WT | This study |
| | *ΔtssM124tssB3* | ΔT6SS; in-frame deletion of *LYSZa7_RS00480, LYSZa7_RS14210, LYSZa7_RS08600 and LYSZa7_RS17780* in LYSZa7 WT | This study |
| | *ΔtssM12tssB3 Para-trc* | H4-Para; replacement of the H4 promoter to ParaBAD-trc promoter in LYSZa7 H4 | This study |
| | *ΔtssM124tssB3 Para-trc* | ΔT6SS-Para; replacement of the H4 promoter to ParaBAD-trc promoter in LYSZa7 ΔT6SS | This study |
| | *tssB4_sfGFP* | Chromosomal C-terminal fusion of the sfGFP to TssB4 | This study |
| | *tssH4_sfGFP* | Chromosomal C-terminal fusion of the sfGFP to TssH4 | This study |
| | *ΔtssM12tssB3 tssB4_sfGFP* | H4 tssB4_sfGFP; chromosomal C-terminal fusion of the sfGFP to TssB4 in LYSZa7 H4 | This study |
| | *ΔtssM124tssB3 tssB4_sfGFP* | ΔT6SS tssB4_sfGFP; chromosomal C-terminal fusion of the sfGFP to TssB4 in LYSZa7 ΔT6SS | This study |
| | *ΔtssM12tssB3 tssH4_sfGFP* | H4 tssH4_sfGFP; chromosomal C-terminal fusion of the sfGFP to TssH4 in LYSZa7 H4 | This study |
| | *ΔtssM124tssB3 tssH4_sfGFP* | tssH4_sfGFP; chromosomal C-terminal fusion of the sfGFP to TssH4 in LYSZa7 ΔT6SS | This study |
| | *tssB4_sfGFP Para-trc* | WT-Para tssB4_sfGFP; replacement of the H4 promoter to ParaBAD-trc promoter in LYSZa7 *tssB4_sfGFP* | This study |
| | *ΔtssM12tssB3 tssB4_sfGFP Para-trc* | H4-Para tssB4_sfGFP; replacement of the H4 promoter to ParaBAD-trc promoter in LYSZa7 H4 *tssB4_sfGFP* | This study |
| | *ΔtssM124tssB3 tssB4_sfGFP Para-trc* | ΔT6SS-Para tssB4_sfGFP; replacement of the H4 promoter to ParaBAD-trc promoter in LYSZa7 ΔT6SS *tssB4_sfGFP* | This study |
| | *tssB4_sfGFP Para-trc Δhcp4* | In-frame deletion of LYSZa7_RS08500 in LYSZa7 WT-Para | This study |
| | *ΔtssM12tssB3 tssB4_sfGFP Para-trc Δhcp4* | In-frame deletion of LYSZa7_RS08500 in LYSZa7 H4-Para | This study |
| | *ΔtssM124tssB3 tssB4_sfGFP Para-trc Δhcp4* | In-frame deletion of LYSZa7_RS08500 in LYSZa7 ΔT6SS-Para | This study |
| | *ΔtssM12tssB3 tssB4_sfGFP Para-left* | H4-Para-left; replacement of the *tssB4* promoter to ParaBAD promoter in LYSZa7 H4 *tssB4_sfGFP* | This study |
| | *ΔtssM12tssB3 tssB4_sfGFP Para-right* | H4-Para-right; replacement of the *tssH4* promoter to ParaBAD promoter in LYSZa7 H4 *tssB4_sfGFP* | This study |
| | *ΔtssM12tssB3 tssB4_sfGFP Para-trc Δ8480* | In-frame deletion of LYSZa7_RS08480 in LYSZa7 H4-Para | This study |
| | *ΔtssM12tssB3 tssB4_sfGFP Para-trc Δ8520* | In-frame deletion of LYSZa7_RS08520 in LYSZa7 H4-Para | This study |
| | *ΔtssM12tssB3 tssB4_sfGFP Para-trc Δ8480Δ8520* | In-frame deletion of LYSZa7_RS08480 and LYSZa7_RS08520 in LYSZa7 H4-Para | This study |
| | *ΔtssM12tssB3 Δ8480-8740* | In-frame deletion of LYSZa7_RS08480-RS08470 in LYSZa7 H4 | This study |
| | *ΔtssM12tssB3 Δ8520-8515* | In-frame deletion of LYSZa7_RS08520-RS08515 in LYSZa7 H4 | This study |
| | *ΔtssM12tssB3 tssH4_sfGFP Para-trc ΔvgrG4b* | In-frame deletion of LYSZa7_RS08530 in LYSZa7 H4-Para tssH4_sfGFP | This study |
| | *ΔtssM12tssB3 tssH4_sfGFP Para-trc Δpaar4* | In-frame deletion of LYSZa7_RS08505 in LYSZa7 H4-Para tssH4_sfGFP | This study |
| | *ΔtssM12tssB3 tssH4_sfGFP Para-trc ΔDUF4123* | In-frame deletion of LYSZa7_RS08525 in LYSZa7 H4-Para tssB4_sfGFP | This study |
| | *ΔtssM12tssB3 tssB4_sfGFP ΔmvaT* | In-frame deletion of LYSZa7_RS07875 in LYSZa7 H4 tssB4_sfGFP | This study |
| | *ΔtssM12tssB3 tssB4_sfGFP ΔmvaU* | In-frame deletion of LYSZa7_RS19305 in LYSZa7 H4 tssB4_sfGFP | This study |
| | *ΔtssM12tssB3 tssB4_sfGFP ΔmvaU-mvaT77** | Truncationin allele of LYSZa7_RS07875 at residue 77 with a premature stop codon in LYSZa7 H4 ΔmvaU tssB4_sfGFP | This study |
| | *ΔtssM12tssB3 tssH4_sfGFP ΔmvaT* | In-frame deletion of LYSZa7_RS07875 in LYSZa7 H4 tssH4_sfGFP | This study |
| | *ΔtssM12tssB3 tssH4_sfGFP ΔmvaU* | In-frame deletion of LYSZa7_RS19305 in LYSZa7 H4 tssH4_sfGFP | This study |
| | *ΔtssM12tssB3 tssH4_sfGFP ΔmvaU-mvaT77** | Truncationin allele of LYSZa7_RS07875 at residue 77 with a premature stop codon in LYSZa7 H4 ΔmvaU tssH4_sfGFP | This study |

| | | | |
|---|---|---|---|
| | *ΔtssM12tssB3 Δsfa4* | In-frame deletion of LYSZa7_RS08585 in LYSZa7 H4 | This study |
| | ΔtssM12tssB3 Para-trc Δsfa4 | In-frame deletion of LYSZa7_RS08585 in LYSZa7 H4 Para | This study |
| ***Pseudomonas aeruginosa*** **PAO1** | WT | Strain used for T6SS killing and expressing TssB2-sfGFP and TssB4-sfGFP | Lab stock |
| | Δ*tssB123* | Strain used for T6SS killing and expressing TssB2-sfGFP and TssB4-sfGFP | Lab stock |
| | Δ*tssB13* | Strain used for expressing TssB2-sfGFP and TssB4-sfGFP | Lab stock |
| ***Vibrio cholerae*** **V52** | | Prey used for T6SS killing | Lab stock |
| ***Escherichia coli*** **T-Fast** | | Strain used for cloning and gene expression | Lab stock |
| ***E. coli*** **SM10$^{strR}$ λ pir** | streptomycin resistance | Prey used for T6SS killing | Lab stock |
| ***E. coli*** **MG1655** | | Prey used for T6SS killing | Lab stock |
| ***E. coli*** **BL21(DE3)** | | Strain used for protein purification | Lab stock |
| ***E. coli*** **WM6026** | | Strain used for conjugation | Lab stock |

| **plasmid** | **Source** |
|---|---|
| pEXG2.0 | Lab stock |
| pEXG2.0-Δ*tssM1* | This study |
| pEXG2.0-*ΔtssM2* | This study |
| pEXG2.0-*ΔtssB3* | This study |
| pEXG2.0-*ΔtssM4* | This study |
| pEXG2.0-*tssB4-sfGFP* | This study |
| pEXG2.0-*tssH4-sfGFP* | This study |
| pEXG2.0-*Para-trc* | This study |
| pEXG2.0-*Δhcp4* | This study |
| pEXG2.0-*Δsfa4* | This study |
| pEXG2.0-*Δ8480* | This study |
| pEXG2.0-*Δ8480-8470* | This study |
| pEXG2.0-*Δ8520* | This study |
| pEXG2.0-*Δ8520-8515* | This study |
| pEXG2.0-*ΔvgrG4b* | This study |
| pEXG2.0-*Δpaar4* | This study |
| pEXG2.0-*ΔDUF4123* | This study |
| pEXG2.0-*ΔmvaT* | This study |
| pEXG2.0-*ΔmvaU* | This study |
| pEXG2.0-*mvaT77** | This study |
| pPSV37 | Lab stock |
| pPSV37-Sfa4-Flag | This study |
| pPSV37-TssB4-Flag | This study |
| pPSV37-TssB2-Flag | This study |
| pPSV37-8515-Flag | This study |
| pPSV37-8520-8515-Flag | This study |
| pPSV37-8480-8470-Flag | This study |
| pPSV37-8520ΔCD-Flag | This study |
| pPSV37-8520GYAA-Flag | This study |
| pPSV37-VgrG4b-Flag | This study |
| pPSV37-PAAR4-Flag | This study |
| pPSV37-DUF4123-Flag | This study |
| pPSV37-MvaT-Flag | This study |
| pPSV37-MvaU-Flag | This study |
| pPSV37-MvaT-3V5-ara-MvaU-Flag | This study |
| pET28a-His-Hcp4 | This study |
| pET28a-His-MvaT | This study |
| pET28a-His-8520 | This study |
| pET28a-His-GFP-8520 | This study |
| pBAD24 | Lab stock |
| pBAD24-Tat | This study |
| pBAD24-8520-3V5 | This study |
| pBAD24-Tat-8520-3V5 | This study |
| pBAD24-Tat-8520-8515-3V5 | This study |
| pBAD24-Tat-8520ΔTMD1-3V5 | This study |
| pBAD24-Tat-8520ΔTMD2-3V5 | This study |
| pBAD24-Tat-8520ΔTMD3-3V5 | This study |
| pBAD24-Tat-8520ΔTMD4-3V5 | This study |
| pBAD24-8480-3V5 | This study |
| pBAD24-8480-8475-3V5 | This study |
| pBAD24-8480-8475-8470-3V5 | This study |

| | |
|---|---|
| pBAD24-8480K205-216A-3V5 | This study |
| pBAD24-8480K224-240A-3V5 | This study |
| pBAD24-8480K246-254A-3V5 | This study |
| pBAD24-8480K257-266A-3V5 | This study |
| pBAD24-8480K271-278A-3V5 | This study |
| pBAD24-8480K287-298A-3V5 | This study |
| pBAD24-8480K301-308A-3V5 | This study |
| pPSV37Δlac-PtssB4-lux | This study |
| pPSV37Δlac-PtssH4-lux | This study |
| pPSV37Δlac-Phcp4-lux | This study |
| pPSV37Δlac-PtssB4-sfGFP | This study |
| pPSV37Δlac-PtssH4-sfGFP | This study |
| pPSV37Δlac-Phcp4-sfGFP | This study |

**Supplementary Table S2. cyro-EM_dataprocess.**

| | |
|---|---|
| Data Set | RS08520 |
| PDB | 22RM |
| EMDB | 68622 |
| **Data collection and processing** | |
| Micrographs | 8704 |
| Microscope | Titan Krios |
| Voltage(KV) | 300 |
| Electron Exposure(e/Å$^2$) | 50 |
| Defocus Range(μm) | -1.0~-2.0 |
| Pixel size(Å) | 0.46 |
| Initial particle images | 1,613,201 |
| Final particle images | 305,916 |
| Map resolution(A) | 3.0 |
| FSC threshold | 0.143 |
| Map sharpening B factor | 153 |
| **Refinement** | |
| **Model composition** | |
| Non-hydrogen atoms | 5791 |
| Protein residues | 749 |
| **B-factors** | |
| protein | 65.66 |
| **RMS deviations** | |
| Bond length(Å) | 0.002 |
| Bond angle(degree) | 0.457 |
| **Validation** | |
| Mol probity score | 1.77 |
| Clashscore | 3.28 |
| Poor rotamers(%) | 2.87 |
| **Ramachadran** | |
| Favored(%) | 95.62 |
| Allowed(%) | 4.38 |
| Outliers(%) | 0 |
