## Supplementary material for "Functional activation reveals a repressor-gated fourth type VI secretion system in clinical *Pseudomonas aeruginosa*": Fig. S1-11

Liwen Wu *et al.*

* Tao Dong.

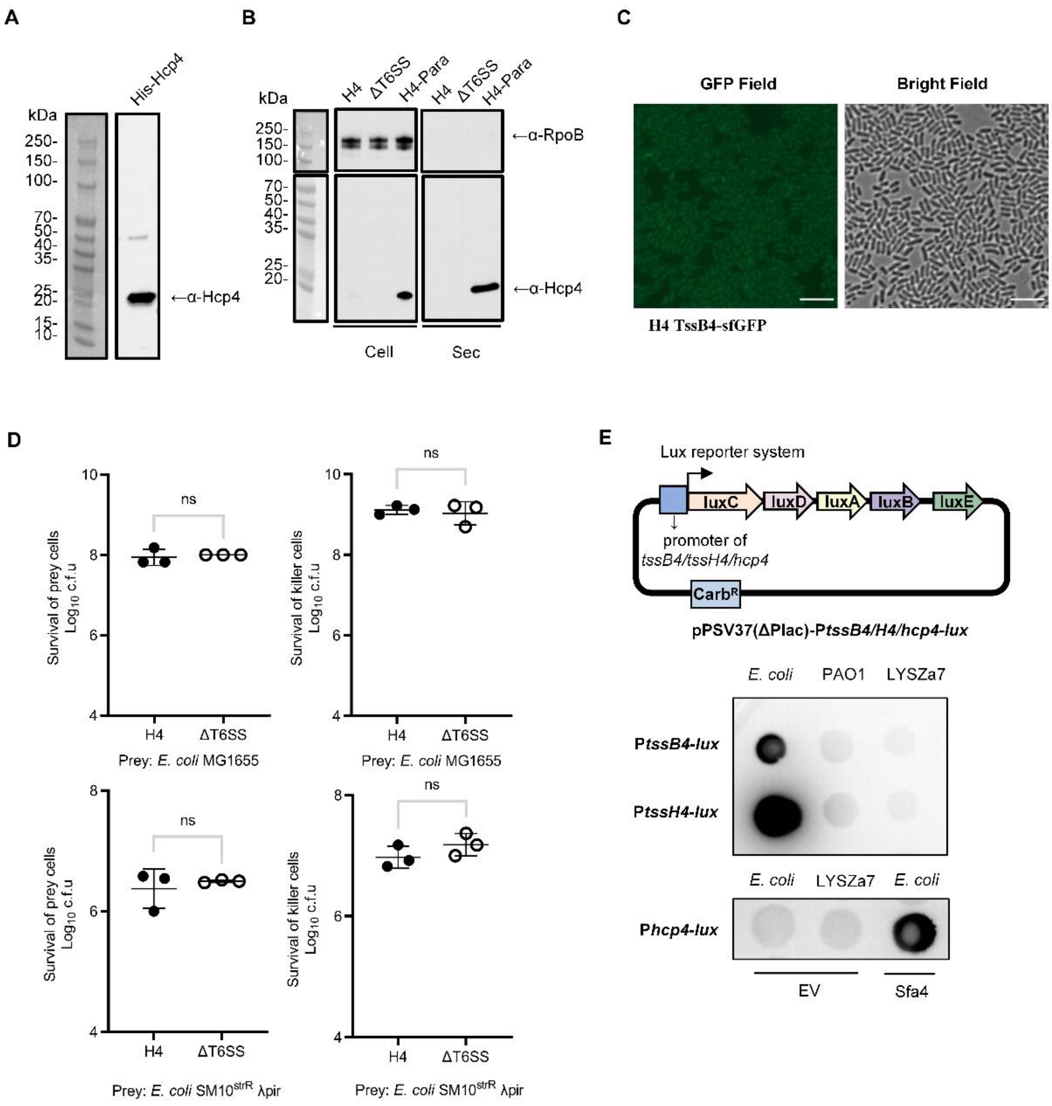


**Supplementary Figure 1. H4-T6SS is silenced under laboratory conditions.**

**A.** Hcp4 antibody detection. The molecular weight of the purified His-Hcp4 protein is 21.27 kDa. The concentration of the protein sample is 0.1 µg. **B.** Hcp4 expression and secretion levels in H4 (Δ*tssM1tssM2tssB3*) and ΔT6SS (Δ*tssM1tssM2tssB3tssM4*) strains. The induced H4-Para strain was used as a positive control. Cells were grown at 37°C for 4 h to $OD_{600}$ ~1. **C.** Fluorescence images of sfGFP-labeled TssB4 in H4 strain. Cells were grown at 28°C for 6 h to $OD_{600}$ ~1. Scale bar: 5 µm. **D.** Competition assays of H4 and Δ T6SS against *E. coli* MG1655 and SM10$^{strR}$ λpir. Cells of killer and prey were mixed at a ratio of 10:1 (killer: prey), and co-incubated for 12 h at 30 °C. Error bars indicate the standard deviation of three biological replicates and statistical significance was calculated using a Student's t-test for each group, ns, not significant. **E.** Expression of H4-T6SS promoters. The original inducible lacUV5 promoter of the pPSV37 vector was replaced with the *tssB4*, *tssH4* or *hcp4* promoter, and luciferase reporter genes were fused. *E. coli* or *P. aeruginosa* cells carrying a report plasmid were cultured to $OD_{600}$ ~1 and were then plated on the LB-agar plate. Cells carrying additional empty (EV) or Sfa4 expression plasmid were plated on the LB-agar plate containing 1 mM IPTG. After incubation at 37°C for 6 h, the signal was detected by chemiluminescence.

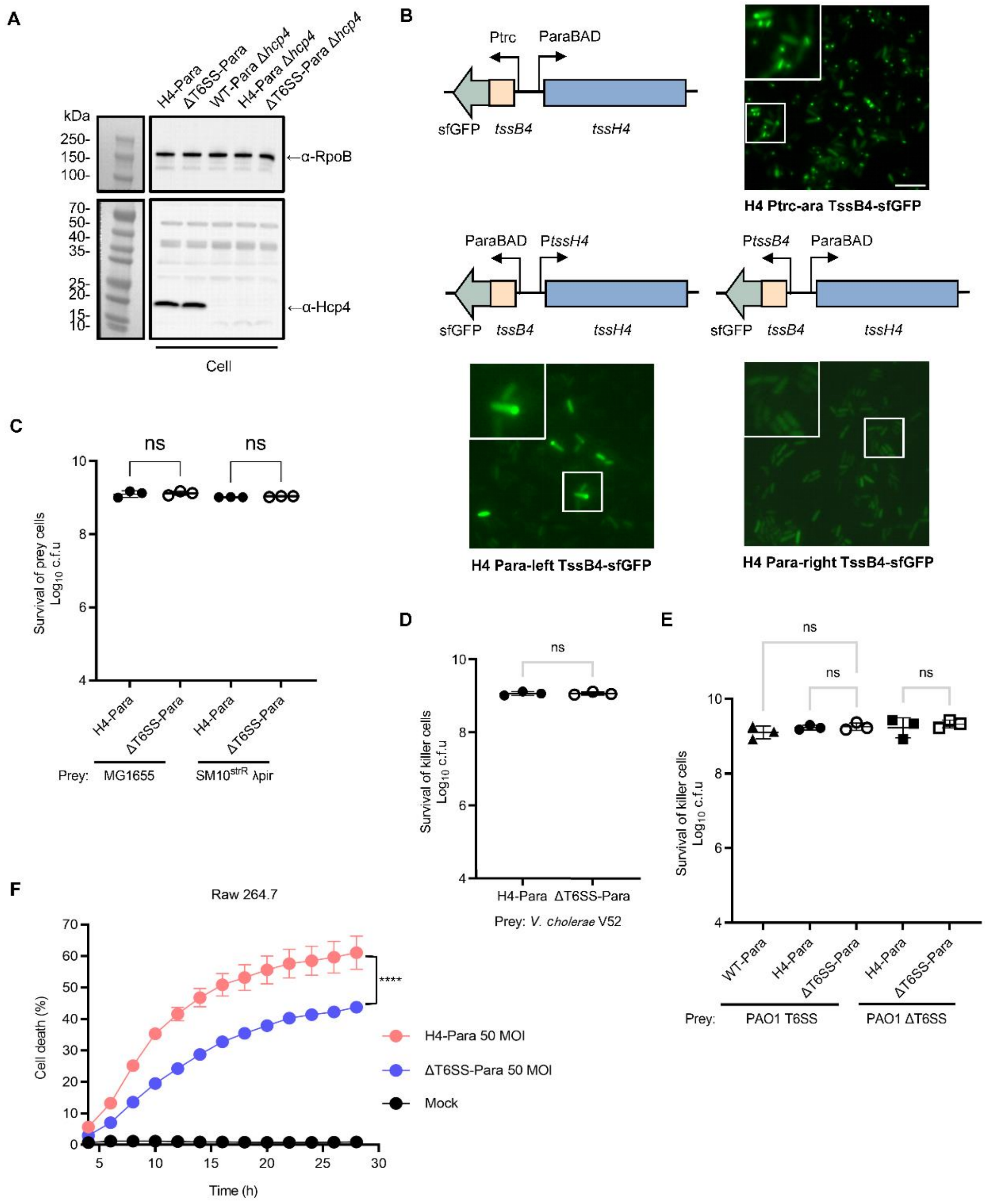


**Supplementary Figure 2. Induced H4-T6SS exhibits dynamic assembly and competitive advantage of T6SS.**

**A.** Detection of Hcp4 antibody specificity. **B.** Establishment of activation conditions for H4-T6SS. Inducible promoter was used to replace either both arms of the promoters or only one of the promoters. Fluorescence images showing sheath assembly in H4 Ptrc-ara but not in H4 Para-left or Para-right strains. Cells were induced with 0.1% [w/v] arabinose for 3 h during the exponential phase. A representative 30 × 30 μm field of cells with a 3× magnified 5 × 5 μm inset (marked by box) is shown. Scale bar: 5 μm. **C-E.** Survival of killer strains in the competition assays depicted in Figure 1E-G. Error bars indicate the standard deviation of three biological replicates and statistical significance was calculated using a Student's t-test (**C** and **D**) and a one-way ANOVA test (**E**) for each group, ns, not significant. **F.** Cytotoxicity detection of H4-Para and Δ T6SS-Para strains to Raw254.7 cell. Host cells were incubated with the bacteria cells (MOI 50) for 2 h. 0.2 μg/mL propidium iodide (PI) was added to the media for IncuCyte experiments. Statistical significance was calculated using a two-way ANOVA test, ns, not significant, ****$p < 0.0001$.

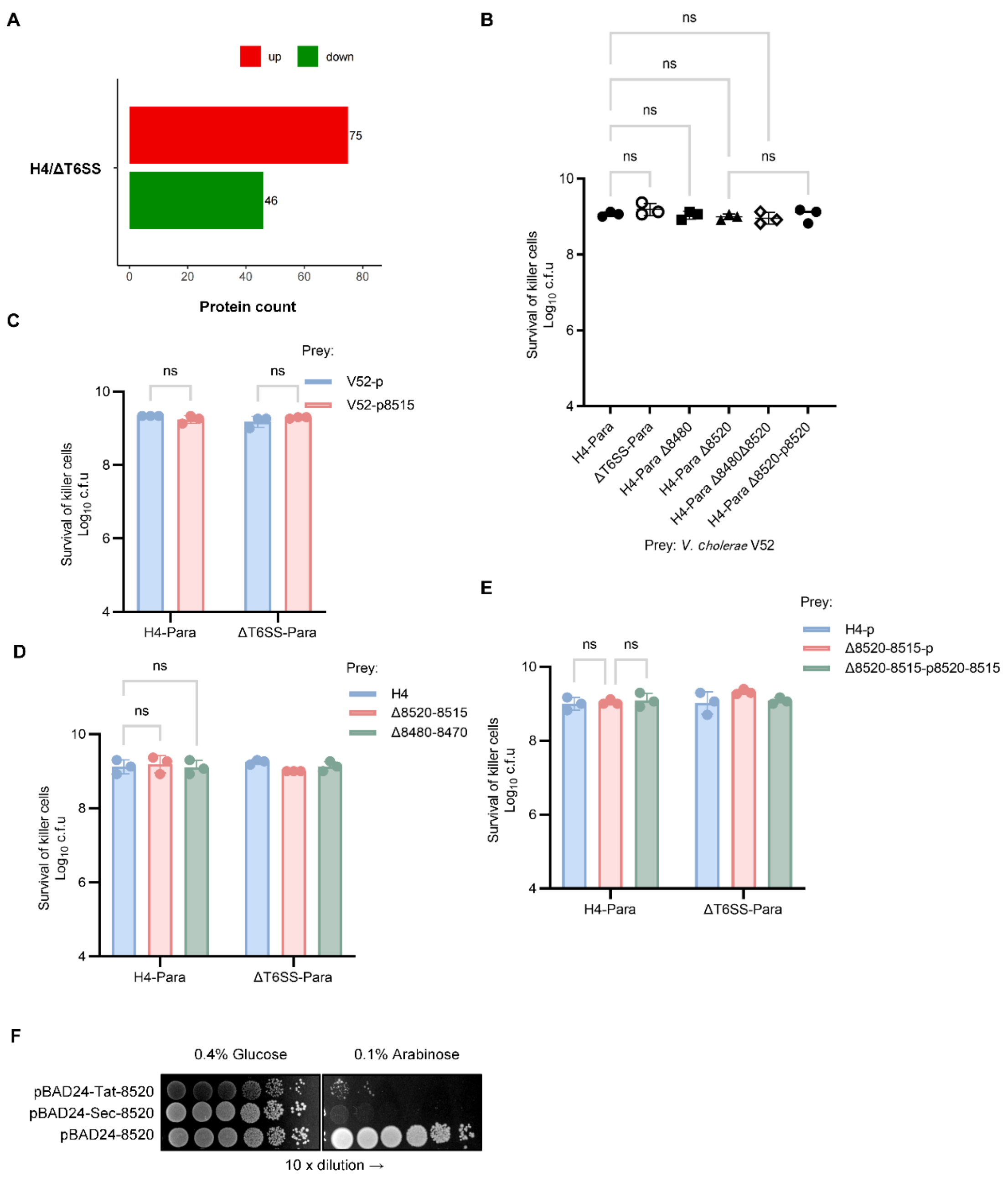


**Supplementary Figure 3. Secretome analysis reveals new effectors of H4-T6SS.**

**A.** Screening of pairwise phenotypic differential proteins. Differentially secreted proteins were screened out by using a Fold change (FC) greater than 1.5 times and t-test statistics to obtain a significance (p-value) less than 0.05. The number of significantly different proteins enriched in the supernatant of H4-Para compared to that of ΔT6SS-Para is shown. **B-E.** Survival of killer strains in the inter- and intraspecific competition assays depicted in Figure 2B-E. Error bars indicate the standard deviation of three biological replicates and statistical significance was calculated using a one-way ANOVA test, ns, not significant. **F.** Bacterial toxicity assay of effector RS08520 when targeting *E. coli* cytoplasm and periplasm. *E. coli* cells were plated on 0.1% [w/v] arabinose and 0.4% [w/v] glucose plates for protein induction and repression, respectively.

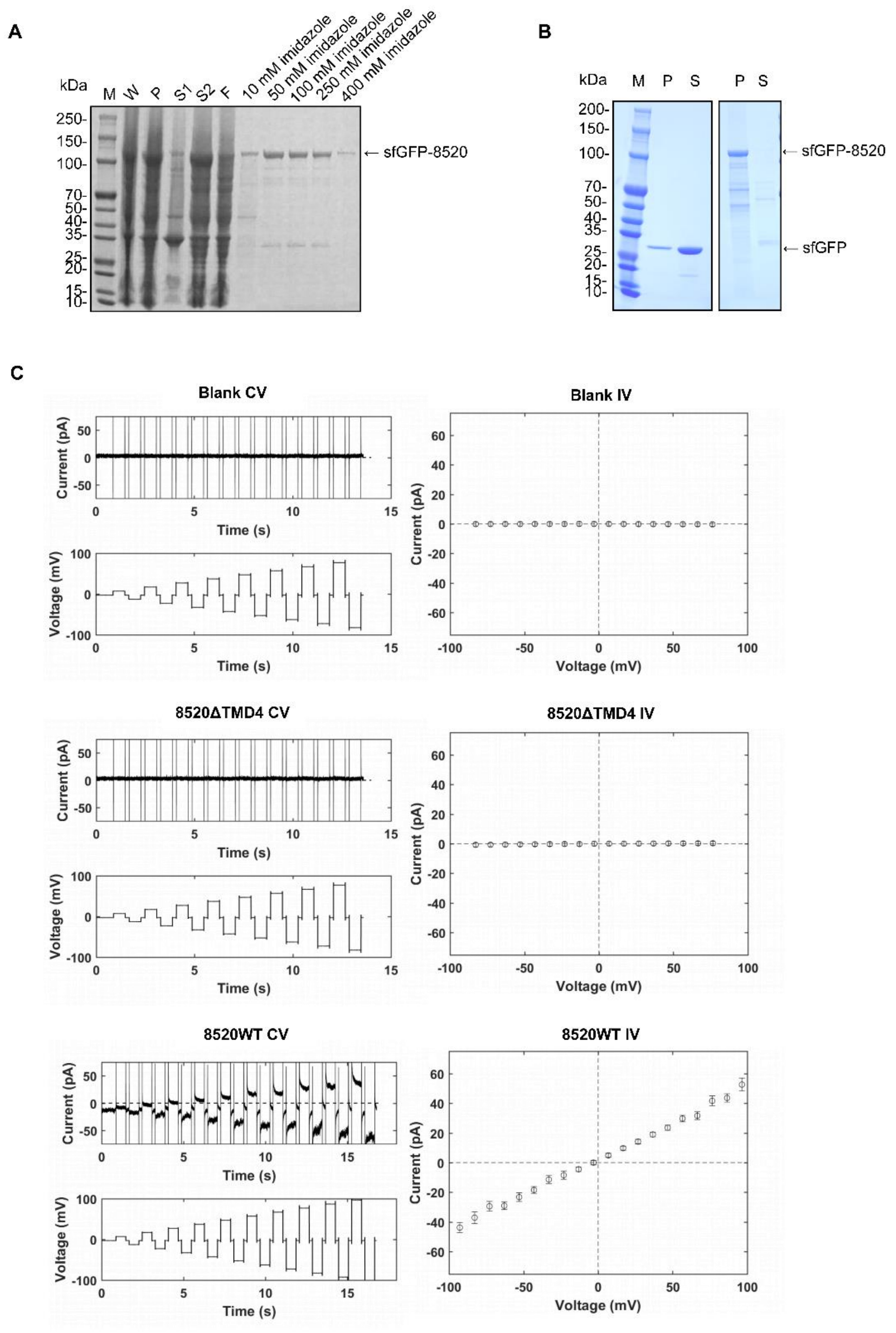


**Supplementary Figure 4. Effector RS08520 exhibits membrane pore-forming activity.**

**A.** SDS-PAGE analysis of the purified sfGFP-RS08520 protein. M: protein marker; W: the whole-cell sample; P: the pellet sample of the cell lysate; S1: the supernatant sample of the cell lysate after centrifugation at 3,000 ×*g*; S2: the supernatant sample of S1 after centrifugation at 15,000 ×*g*; F: the flow through of the Ni-NTA. **B.** Detection of the liposomes binding property of RS08520 protein *in vitro*. S: the supernatant sample; P: the pellet of liposome sample. **C.** Detection of the membrane pore-forming characteristics of the effector RS08520. After adding 0.2 μM purified RS08520 and TMD-mutated proteins to the DPhPC membrane for 5 min, a stable current trace was formed as shown. The applied voltages of each sample are shown at the bottom. I/V curves corresponding to the current traces are shown in the right panel.

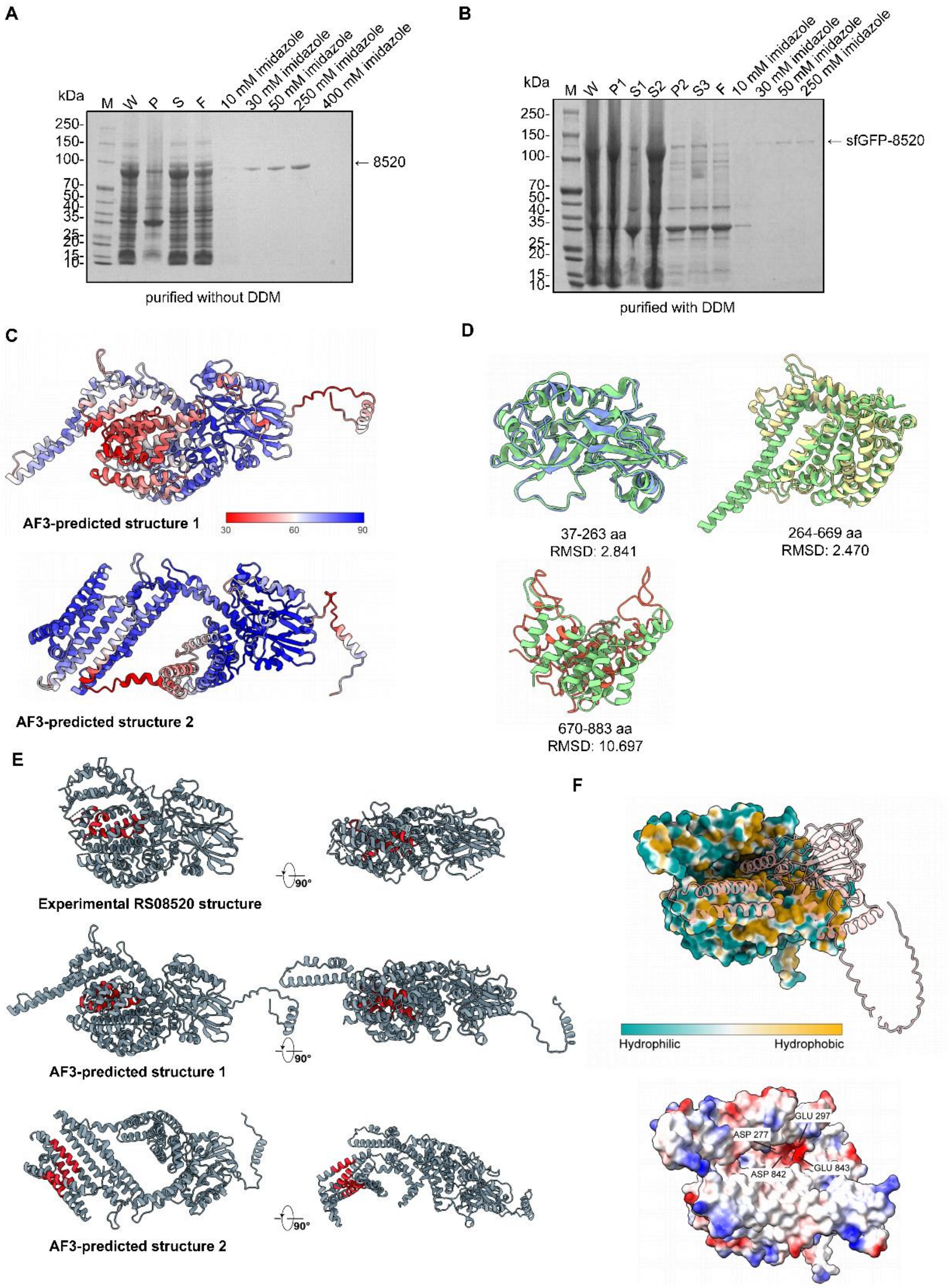


**Supplementary Figure 5. Structural characterization of the effector RS08520.**

**A.** SDS-PAGE analysis of the purified RS08520 protein. M: protein marker; W: the whole-cell sample; P: the pellet sample of the cell lysate; S: the supernatant sample of the cell lysate; F: the flow through of the Ni-NT. **B.** SDS-PAGE analysis of the sfGFP-

RS08520 protein purified from cell membrane. M: protein marker; W: the whole-cell sample; P1: the pellet sample of the cell lysate at 3,000 ×*g*; S1: the supernatant sample of the cell lysate at 3,000 ×*g*; P2: the pellet sample of S1 (containing cell membrane) at 15,000 ×*g*; S2: the supernatant sample obtained by resuspending P2 in the lysis buffer containing 1% DDM and rotating for 1 h; S3: the supernatant sample of S2 at 15,000 ×*g*. F: the flow through of the Ni-NTA. **C.** AlphaFold 3 predicted models of RS08520 colored by pLDDT confidence scores. Two different protein conformations of RS08520 is shown. **D.** Structural superposition of the experimental RS08520 structure and the AlphaFold 3 predicted model 1. NTD, the central linker and CTD are colored in blue, yellow and red, respectively. AlphaFold 3 predicted model is colored in green. **E.** Location of predicted transmembrane helices (colored red) mapped onto the structural models. **F.** AlphaFold 3-predicted complex of RS08520 and its immunity protein. The RS08520 surface is colored by hydrophobicity, while the immunity protein is shown in ribbon representation (upper panel). The RS08520 surface is colored by electrostatic potential (lower panel).

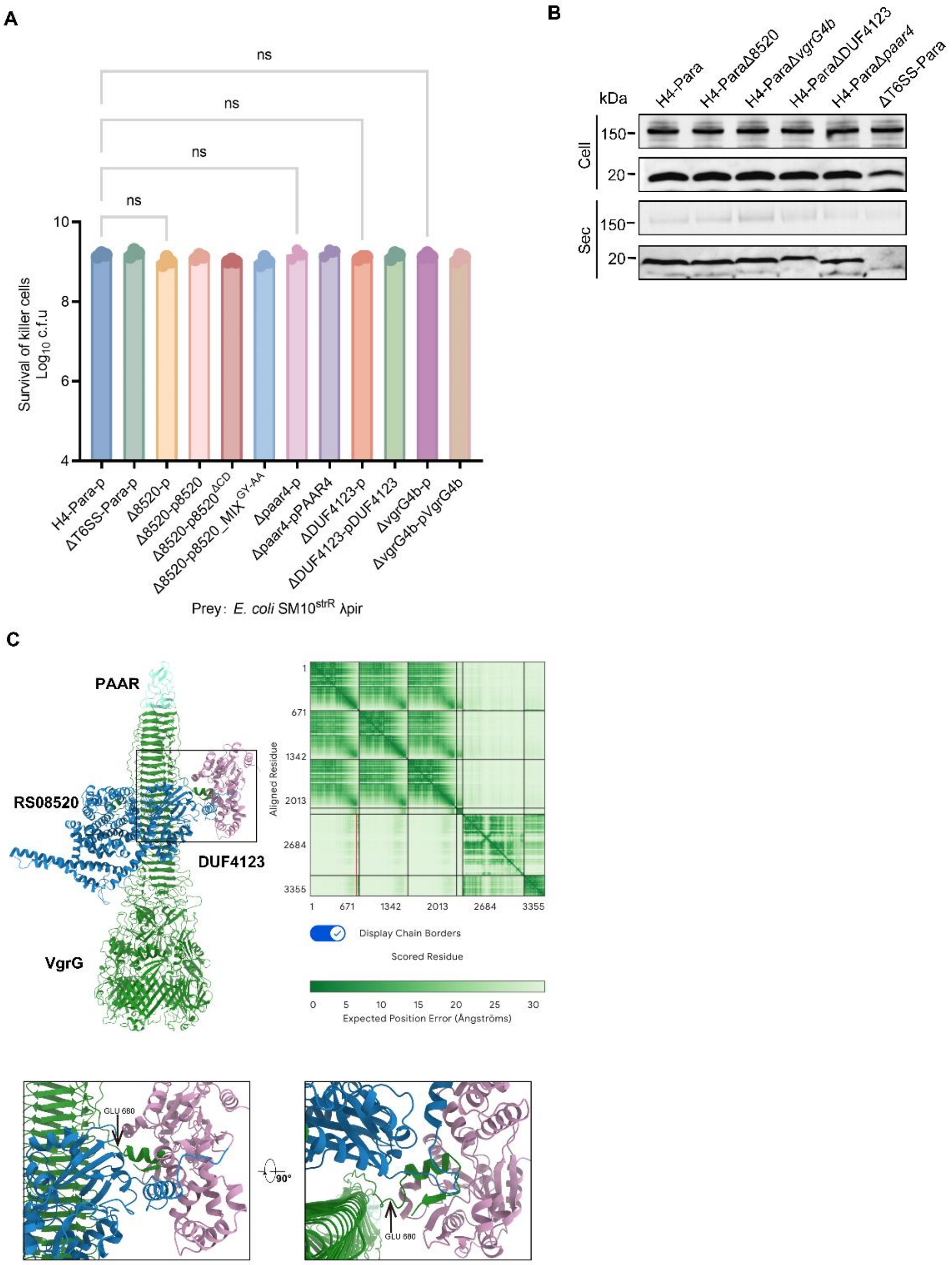


**Supplementary Figure 6. Chaperone–VgrG–PAAR pathway mediates the secretion of RS08520.**

**A.** Survival of killer strains in the competition assay depicted in Figure 3E. Error bars indicate the standard deviation of three biological replicates and statistical significance was calculated using a one-way ANOVA test, ns, not significant. **B.** Effects of the absence of RS08520 and its upstream and downstream proteins on the secretion of H4-Para. **C.** AlphaFold 3-predicted complex comprising RS08520, VgrG4b, PAAR4, and the DUF4123 chaperone, alongside the Predicted Aligned Error (PAE) plot for the complex. Side and bottom-up views of the RS08520 loading site are shown as a zoom-in of the boxed region.

A

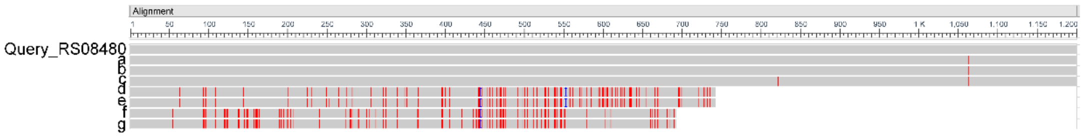


| Description | Query Cover | E value | Per. Ident |
|---|---|---|---|
| Pseudomonas aeruginosa strain LYSZa7 | 100% | 0.0 | 100.00% |
| a Pseudomonas aeruginosa strain ZYPA192 | 100% | 0.0 | 99.92% |
| b Pseudomonas aeruginosa strain ZYPA162 | 100% | 0.0 | 99.92% |
| c Pseudomonas aeruginosa MTB-1 | 100% | 0.0 | 99.83% |
| d Pseudomonas aeruginosa strain CF-sbj13-Pa1-I | 62% | 0.0 | 88.02% |
| e Pseudomonas aeruginosa strain CF-sbj14-Pa1-I | 62% | 0.0 | 87.89% |
| f Pseudomonas aeruginosa strain PaLo22 | 58% | 0.0 | 86.87% |
| g Pseudomonas aeruginosa strain Paer81 | 58% | 0.0 | 86.87% |

B

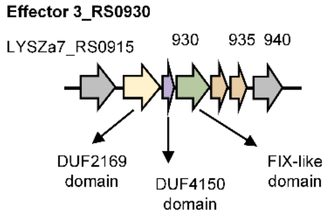


**Supplementary Figure 7. Effector RS08480 is relatively conserved in *P. aeruginosa* H4-T6SS positive strains.**

**A.** Alignment of the RS08480 nucleotide sequence in LYSZa7 strain with the homologous genes in other H4-T6SS positive strains. BLAST identified 7 genes of the *P. aeruginosa* that have a high similarity to RS08480. Mismatches are shown in red. **B.** Schematic diagram of effector RS0930 gene island. Bioinformatics analysis predicts that RS0930 and RS08480 have a similar functional domain.

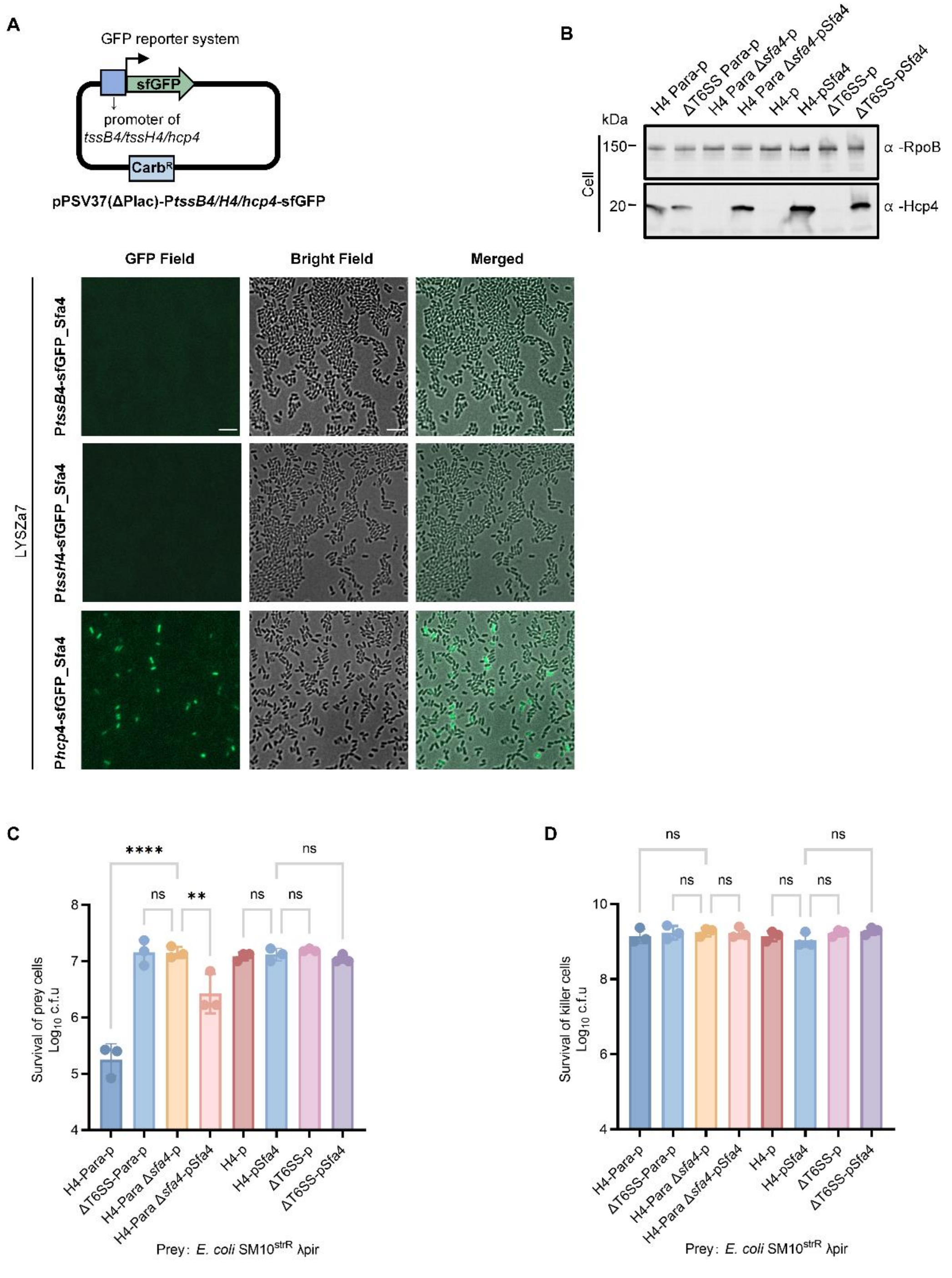


**Supplementary Figure 8. Sfa4 regulator is required for *hcp4* expression.**

**A.** Detection of H4-T6SS promoter expression. The original inducible lacUV5 promoter of the pPSV37 vector was replaced with the *tssB4*, *tssH4* or *hcp4* promoter, and *sfgfp* reporter gene was fused. Fluorescence images showing the activation of Sfa4 on the H4-promoter reporter plasmids in LYSZa7. Scale bar: 5 μm. Cells carrying a report plasmid were cultured to $OD_{600}$ ~1 in LB medium with 1 mM IPTG. **B.** Expression of Hcp4 in H4-Para and H4 strains with Sfa4 deficiency and overexpression. Cells were grown at 37°C for 4 h to $OD_{600}$ ~1, with 0.1% [w/v] arabinose and 1 mM IPTG. **C.** Competition assay of the H4-Para, ΔT6SS-Para, H4, ΔT6SS and Sfa4 deficiency or overexpression mutants. Cells of killer and prey were mixed at a ratio of 10:1 (killer: prey), and co-incubated for 12 h at 30°C. During the liquid culture and co-incubation process, 0.1% [w/v] arabinose and 1 mM IPTG were added. Error bars indicate the standard deviation of three biological replicates and statistical significance was calculated using a one-way ANOVA test, $^{**}P < 0.01$, $^{****}P < 0.0001$, ns, not significant.

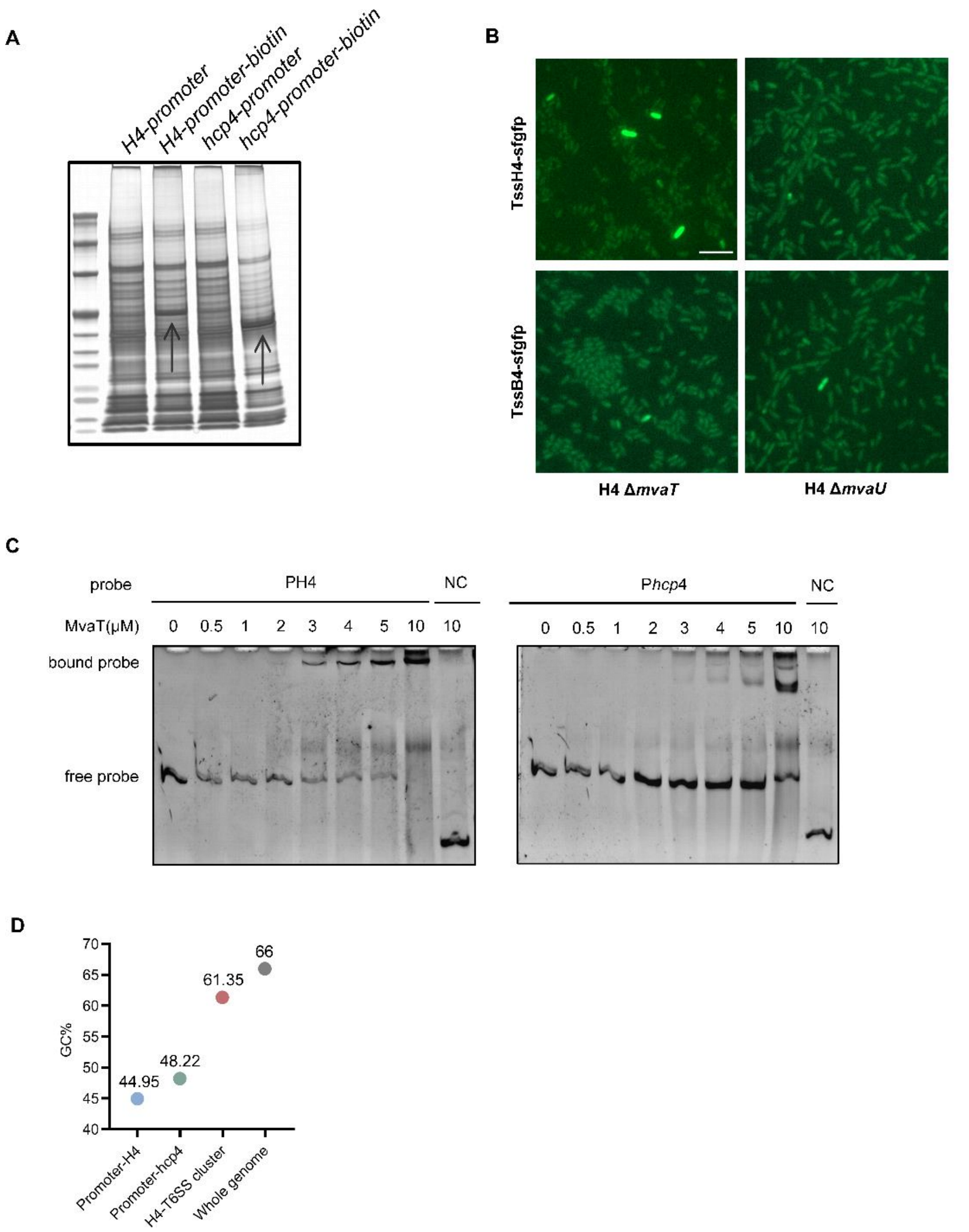


**Supplementary Figure 9. MvaT binds to the H4-T6SS promoter.**

**A.** DNA pull-down assay screening proteins that could specifically bind to the biotinylated H4 and *hcp4* promoters. Arrows indicates possible differential proteins bands. Cell lysates were incubated with each probe, captured with streptavidin beads and analyzed by SDS-PAGE and silver staining. **B.** Fluorescence images showing sfGFP signals of *tssB4* and *tssH4* in H4 Δ*mvaT* and H4 Δ*mvaU* mutants. A representative 30 × 30 μm field is shown. Scale bar: 5 μm. **C.** EMSA demonstrating the binding of the MvaT protein to the H4 and *hcp4* promoters. The *algD* promoter was used as a negative control (NC). **D.** GC content (%) of the complete genome of the LYSZa7 strain, as well as its H4-T6SS cluster and promoters.

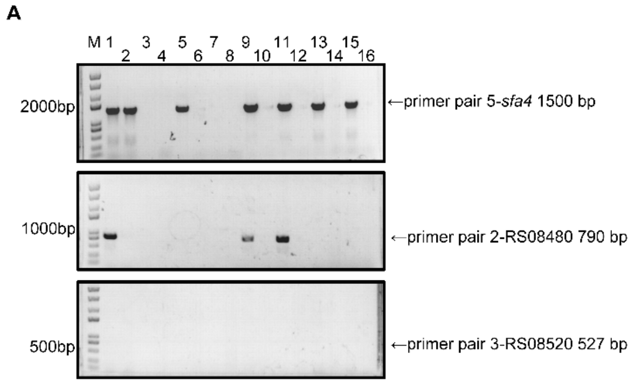


**Supplementary Figure 10. H4-T6SS is prevalent in *P. aeruginosa* clinical isolates.**

**A.** Detection of the H4 genes in *P. aeruginosa* clinical isolates. The nucleic acid gel shows the PCR results of *sfa4*, RS08480 and RS08520 primer pairs in 16 out of ~1200 *P. aeruginosa* strains. Among the 7 strains that could be detected by the *sfa4* primer pair, only 3 could be amplified by the RS08480 primer pair, and no strain could be amplified by the RS08520 primer pair.

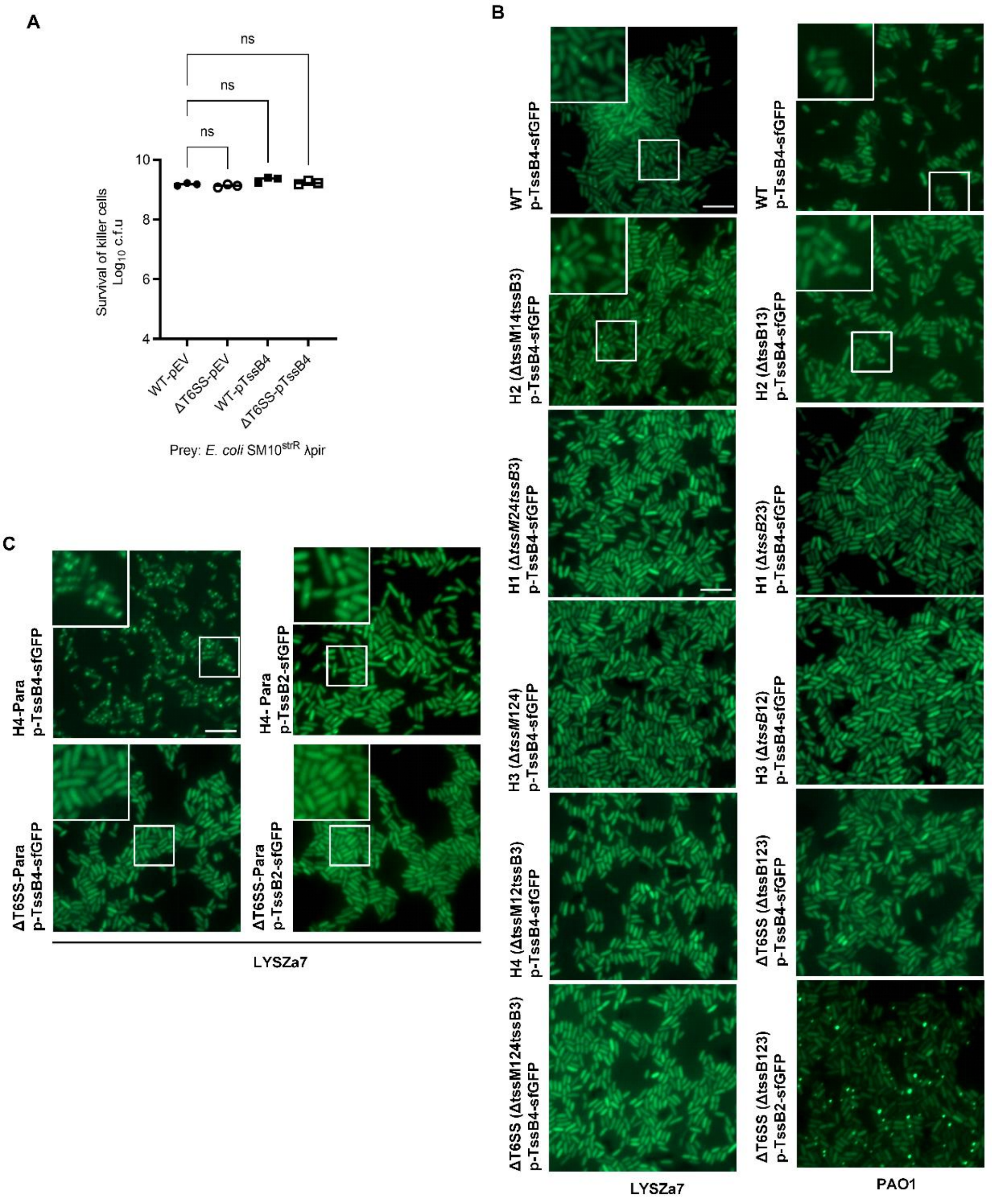


**Supplementary Figure 11. TssB2 is involved in the assembly of H4-T6SS.**

**A.** Survival of killer strains in the competition assay depicted in Figure 7A. Error bars indicate the standard deviation of three biological replicates and statistical significance was calculated using a one-way ANOVA test, ns, not significant. **B.** Fluorescence microscopy images showing that TssB4 is not involved in the assembly of H1- and H3-T6SS. **C.** Fluorescence microscopy images showing the assistance of TssB2 in the assembly of H4-T6SS. For B and C, TssB4 and TssB2 proteins were induced using 1mM IPTG. A representative 30 × 30 μm field of cells with a 3× magnified 5 × 5 μm inset (marked by box) is shown. Scale bar: 5 μm.
